# Genome-wide screen reveals Rab12 GTPase as a critical activator of pathogenic LRRK2 kinase

**DOI:** 10.1101/2023.02.17.529028

**Authors:** Herschel S. Dhekne, Francesca Tonelli, Wondwossen M. Yeshaw, Claire Y. Chiang, Charles Limouse, Ebsy Jaimon, Elena Purlyte, Dario R. Alessi, Suzanne R. Pfeffer

## Abstract

Activating mutations in the Leucine Rich Repeat Kinase 2 (LRRK2) cause Parkinson’s disease. LRRK2 phosphorylates a subset of Rab GTPases, particularly Rab10 and Rab8A, and we showed previously that phosphoRabs play an important role in LRRK2 membrane recruitment and activation (Vides et al., 2022). To learn more about LRRK2 pathway regulation, we carried out an unbiased, CRISPR-based genome-wide screen to identify modifiers of cellular phosphoRab10 levels. A flow cytometry assay was developed to detect changes in phosphoRab10 levels in pools of mouse NIH-3T3 cells harboring unique CRISPR guide sequences. Multiple negative and positive regulators were identified; surprisingly, knockout of the Rab12 gene was especially effective in decreasing phosphoRab10 levels in multiple cell types and knockout mouse tissues. Rab-driven increases in phosphoRab10 were specific for Rab12, LRRK2 dependent and PPM1H phosphatase reversible; they were seen with wild type and pathogenic G2019S and R1441C LRRK2. AlphaFold modeling revealed a novel Rab12 binding site in the LRRK2 Armadillo domain and we show that residues predicted to be essential for Rab12 interaction at this site influence overall phosphoRab levels in a manner distinct from Rab29 activation of LRRK2. Our data support a model in which Rab12 binding to a new site in the LRRK2 Armadillo domain activates LRRK2 kinase for Rab phosphorylation and could serve as a new therapeutic target for a novel class of LRRK2 inhibitors that do not target the kinase domain.

## Introduction

Activating mutations in the large, multidomain, Leucine Rich Repeat Kinase 2 (LRRK2) cause inherited Parkinson’s disease, and lead to the phosphorylation of a subset of Rab GTPases (Alessi and Sammler, 2018; Pfeffer, 2022), particularly Rab8A and Rab10 (Steger et al., 2016; 2017). Rab GTPases function in all steps of membrane trafficking by binding to specific effector proteins in their GTP-bound states (Pfeffer, 2017); they are well known for linking motor proteins to transport vesicles and facilitating the transport vesicle docking process.

LRRK2 phosphorylates a single threonine or serine residue in substrate Rab GTPase switch II domains, and this modification blocks the ability of Rabs to be activated by their cognate guanine nucleotide exchange factors, recycled by GDI protein, or to bind to their effector proteins (Steger et al., 2016; 2017). Instead, phosphorylated Rabs bind to a new set of phosphoRab effectors that include RILPL1, RILPL2, JIP3, JIP4 and MyoVa proteins (Steger et al., 2017; Waschbüsch et al., 2020; Dhekne et al., 2021). Although only a small percentage of a given Rab protein is LRRK2 phosphorylated at steady state (Ito et al., 2016), binding to phosphoRab effectors has a dominant and powerful effect on cell physiology and can interfere with organelle motility in axons (Boecker et al., 2021), primary ciliogenesis (Dhekne et al., 2018; Sobu et al., 2021; Khan et al., 2021) and centriolar cohesion (Lara Ordonez et al., 2021).

We have identified a feed-forward pathway that recruits LRRK2 to membranes and can hold it there to enhance subsequent Rab GTPase phosphorylation (Vides et al., 2022). Multiple Rab GTPases including Rab8A and Rab29 can bind the LRRK2 Armadillo domain at a site (#1) that includes residues R361, R399, L403 and K439 (see also Zhu et al., 2022). Phosphorylated Rab8A and Rab10 GTPases can bind to a second site in the Armadillo domain that includes K17 and K18 (site #2); sites #1 and #2 can be occupied simultaneously (Vides et al., 2022).

We present here an unbiased, genome-wide CRISPR screen in mouse NIH-3T3 cells undertaken to identify regulators of the LRRK2 pathway. Of the multiple positive and negative hits identified, Rab12 was the most potent regulator of LRRK2 activity, when either depleted from cells or overexpressed. We show further our discovery of a third LRRK2 Rab12 binding site (#3) in the Armadillo domain that includes residues E240, S244, and I285; site #3 mutations predicted to block Rab12 binding show decreased phosphoRab10 levels, consistent with a critical role for Rab12 in LRRK2 activation.

## RESULTS

The pooled CRISPR screen to identify modulators of LRRK2 activity utilized mouse NIH-3T3 cells in conjunction with the pooled Brie guide RNA (gRNA) mouse library consisting of 78,637 gRNAs targeting 19,674 genes and an extra 1,000 control gRNAs. [A highly detailed protocol can be found here: (<u>dx.doi.org/10.17504/protocols.io.8epv5jr9jl1b/v1</u>)]. Briefly, a pooled “library” of Cas9-expressing cells is first generated, each cell harboring a different gene knock-out. Genes encoding negative regulators of the LRRK2- phosphoRab10 pathway will increase phosphoRab10 staining when knocked out, and genes encoding positive regulators will decrease phosphoRab10 when knocked-out. Fixed cells are stained with an antibody that specifically and sensitively detects phosphoRab10 and then sorted by flow cytometry to separate cells based on phosphoRab10 content. Gene knockouts responsible for changes in phosphoRab10 levels are then identified by genomic sequencing of cells with higher or lower than normal phosphoRab levels.

Fig. 1A shows an example of flow cytometry of anti-phosphoRab10 stained, control mouse 3T3 cells analyzed under baseline conditions (blue) in relation to MLi-2 treated, LRRK2 inhibited cells (green), secondary antibody-only stained cells (black dashed line) or LRRK2 hyperactivated, nigericin-treated 3T3 cells (pink; cf. Kalogeropulou et al., 2020). The flow cytometry resolution of cells with differing phosphoRab10 levels enabled us to collect the highest 7.5% phosphoRab10 signal and lowest 5% signal and compare these enriched cell populations with unsorted cells. Critical to the success of this method is the ability to obtain non-clumped cells after antibody fixation; otherwise, the average fluorescence of clumps will obscure true hits.

**Figure 1.**
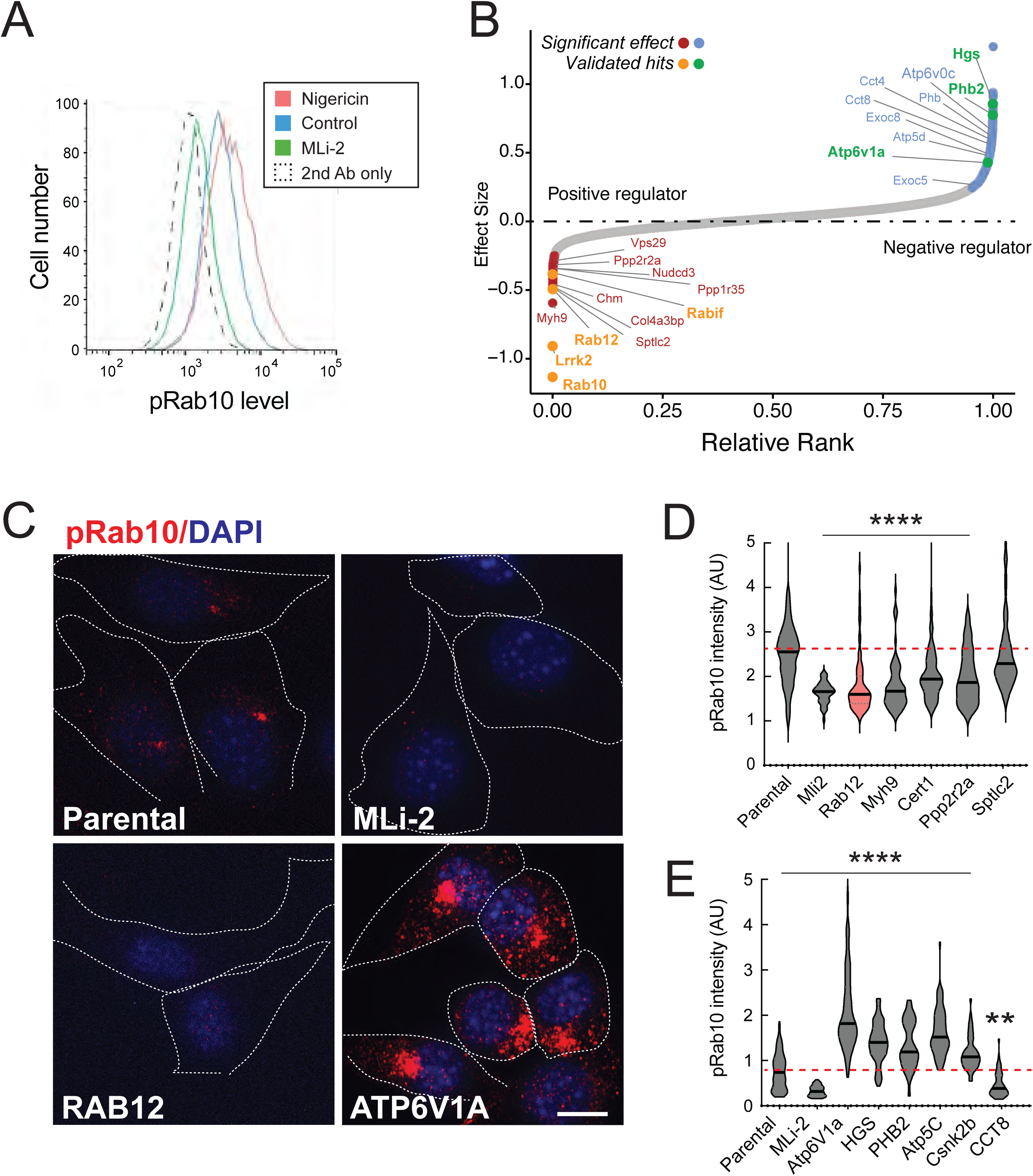
- A flow cytometry based, genome wide CRISPR screen in 3T3-Cas9 cells to reveal modifiers of the LRRK2- phosphoRab10 pathway. (**A)** Phosphorylated Rab10 was detected by flow cytometry after staining cells using anti- phosphoRab10 antibody, either at steady state (control, blue) or in the presence of 4µM Nigericin for 3 h (red) or 200nM MLi-2 for 2 hours. 10,000 cells were analyzed under each of the indicated conditions. **(B)** Statistical analysis of the genome wide screen. After infection with a lentiviral genome-wide CRISPR-Cas9 sgRNA library, genes when knocked-out that reduced (left) or increased (right) phosphoRab10 intensity are indicated on the volcano plot where the X-axis is log2- fold change and Y-axis shows the FDR corrected confidence scores. Genes highlighted are the top positive and negative regulators. **(C,D)** Validation of hits in NIH3T3-Cas9 cells by immunofluorescence microscopy. **(C)** PhosphoRab10was detected by immunofluorescence microscopy in early passage 3T3-Cas9 cells that express lentivirus transduced sgRNAs against the indicated gene after 3 days of Puromycin selection. Scale bar = 10µm. **(D,E)** quantitation of phosphoRab10 fluorescence in cells in which the indicated genes are knocked out. P values: ****, <0.0001; **, 0.0088; n>100 cells counted in two independent experiments.

Statistical analysis of sequencing data from the cells with the lowest phosphoRab10 signal confirmed the success of the screen in that loss of LRRK2, Rab10, and the RABIF Rab10 chaperone (Gulbranson et al., 2017) had the most significant impact on phosphoRab10 expression, as would be expected (Fig. 1B and Figure 1--Figure Supplement 1). Similarly, loss of the CHM gene that is needed for Rab prenylation also led to decreased phosphoRab10. Independent re-validation of the most significant top hits in 3T3 cells (Fig. 1C-E and Fig. 1–Figure Supplement 2) by creating individually knocked out cell lines confirmed most of them, and as will be described below, revealed an unexpected role for Rab12 GTPase.

In addition to Rab12, knockout of genes including MYH9, CERT1, SPTLC2, PPP2R2A, PPP1R35 and NUDCD3 also decreased phosphoRab10 intensity by immunofluorescence microscopy, suggesting that the corresponding gene products are also positive regulators of LRRK2 function (Fig. 1B-D; Fig. 1– Figure Supplements 1 and 2). ER-localized SPTLC2 (serine palmitoyl transferase) is the rate limiting enzyme in ceramide synthesis and CERT1 is critical for ceramide transfer from the ER to the Golgi complex. How ceramide synthesis and transport relate to LRRK2 activity will be addressed in future work; chemical inhibition of SPTLC2 with myriocin did not yield a similar phenotype, suggesting that the role of this pathway in phosphoRab10 regulation may be more complex. PPP2R2A was shown previously to similarly influence phosphoRab10 levels in a phosphatome-wide screen to identify phosphoRab10 phosphatases (Berndsen et al., 2019). PPP1R35 was not tested in that screen but like MYH9, it is involved in primary cilia assembly, and their pericentriolar localizations suggest a connection with phosphoRab10 biology. NUDCD3 stabilizes the dynein intermediate chain and is likely important for concentrating phosphoRab10 at the mother centriole. Finally, 14-3-3 proteins such as YWHAE are known to bind LRRK2 via pSer910 and pSer935 (Nichols et al., 2010) and may stabilize LRRK2 protein.

Knockout of several genes hyperactivated LRRK2 activity and phosphoRab10 levels: these include ATP6V1A, ATP6V0C, HGS, PHB2, ATP5C, and CSNK2B (Fig. 1B, C, E; Fig. 1–Figure Supplements 1 and 2). The ATP6 proteins are non-catalytic subunits of the vacuolar ATPase needed for lysosome acidification; their deletion presumably has similar effects as Bafilomycin that greatly increases LRRK2 activity (cf. Wang et al., 2021). HGS is also known as HRS and is part of the ESCRT-0 complex; loss of HRS function interferes with autophagic clearance and causes ER stress (Oshima et al., 2016).

PHB1/2 are an inner mitochondrial membrane mitophagy receptors that are required for Parkin-induced mitophagy in mammalian cells (Wei et al., 2017). Work from Ganley and colleagues has shown an inverse correlation between LRRK2 activity and mitochondrial turnover (Singh et al., 2021). ATP5C1 is part of the mitochondrial ATP synthase complex V; Casein kinase 1 alpha has been shown to phosphorylate LRRK2 (Chia et al., 2014) but a role for casein kinase 2B is not yet clear. As reported previously by many other groups, lysosomal and mitochondrial stress increased phosphoRab10 levels.

### Loss of Rab12 impacts phosphoRab10 generation

Figure 2 compares the levels of endogenous phosphoRab10 and total Rab10 in parental 3T3 cells, parental cells treated with MLi-2 LRRK2 inhibitor, and a pooled 3T3 cell line in which Rab12 has been knocked out. Quantitation of these data confirmed a roughly five-fold decrease in phosphoRab10 levels under these conditions (Fig. 2B). This was entirely unexpected as prior studies on Rab29, a protein that can activate apparent LRRK2 activity under conditions of protein overexpression (cf. Liu et al., 2018; Purlyte et al., 2018), has no consequence on phosphoRab10 levels in a Rab29 mouse knockout model, in any tissue analyzed or derived mouse embryonic fibroblasts (Kalogeropulou et al., 2020).

**Figure 2.**
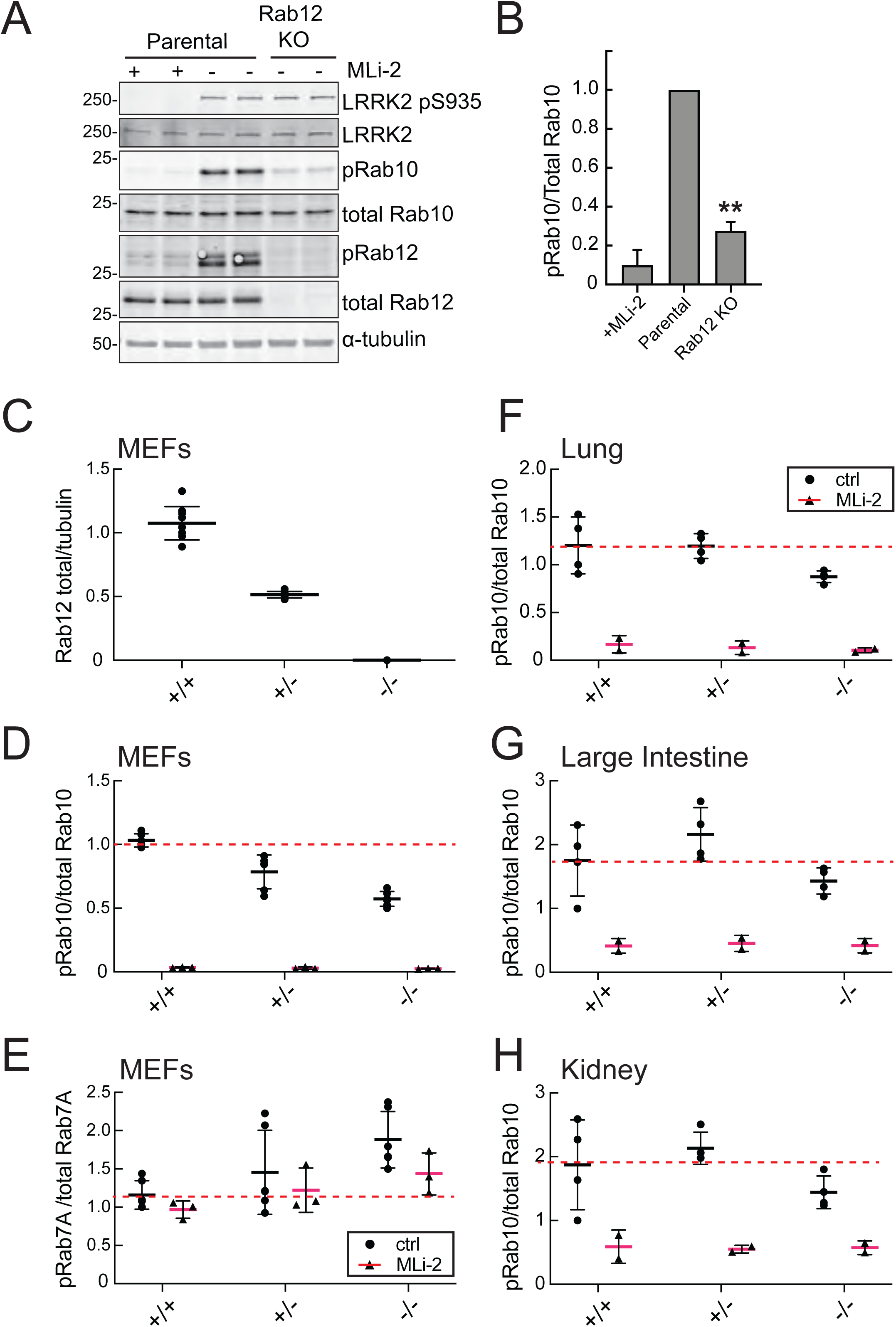
(A,B) Loss of Rab12 decreases phosphoRab10. A) Immunoblot analysis of 3T3-Cas9 cells expressing Rab12 sgRNA (Rab12 KO) or parental cells, +/- MLi2 (200nM for 2h) as indicated. B) Quantitation of phosphoRab10 normalized to total Rab10 from immunoblots in A. Error bars indicate SEM from two experiments carried out in duplicate. **P = 0.002 with Student’s T test. (C-H) Effect of Rab12 knock-out on endogenous LRRK2 activity in embryonic fibroblasts (C-E) and tissues (F-H) derived from Rab12 knockout mice as assessed by immunoblot **(C)** Quantitation of total Rab12 in parental, heterozygous or homozygous knockout Rab12 mice. **(D)** Quantitation of phosphoRab10 normalized to total Rab10 from immunoblots. **(E)** Quantitation of phosphoRab7 normalized to total Rab7 from immunoblots. **(F-G)** Quantitation of phosphoRab10 normalized to total Rab10 from immunoblots of Lung, Large Intestine or Kidney as indicated. MLi-2 was administered to MEFs at 100 nM for 1 h and to mice at 30 mg/kg for 2 h. Blot data is shown in Figure 2-Figure Supplements 1 and 2.

To confirm these data in an animal model, we analyzed cells and tissues derived from Rab12 knock-out mice generated by the Knockout Mouse Phenotyping Program at The Jackson Laboratory using CRISPR technology. Immunoblotting analysis of embryonic fibroblasts (MEFs) confirmed that the heterozygous and homozygous knockouts expressed the expected 50% or 100% loss of Rab12 protein (Fig. 2C; Figure 2–Figure Supplement 1). MEFs derived from homozygous knockout animals showed as much as 50% decrease in phosphoRab10 levels as detected by immunoblot from multiple clones (Fig. 2D); specificity of the detection method was confirmed upon addition of the MLi-2 LRRK2 inhibitor that abolished all phosphoRab10 signal. PhosphoRab7, the product of LRRK1 action (Hanafusa et al., 2019; Malik et al., 2021), appeared to increase moderately as a function of Rab12 loss (Fig. 2E).

Various tissues were analyzed for phosphoRab10 changes in LRRK2 heterozygous and homozygous knockout animals. As shown in Fig. 2 F-H (and Figure 2–Figure Supplement 2), decreases in phosphoRab10 were detected in the homozygous mouse lung with smaller trends in the large intestine and kidney. Together, these data confirm a role for Rab12 in the LRRK2 signaling pathway that is distinct from that of the previously studied Rab29 protein. We were not able to monitor loss of phosphoRab10 in the brain as phosphoRab10 is more difficult to detect in brain tissue that is enriched in the Rab-specific PPM1H phosphatase (Berndsen et al., 2019). Future work will evaluate the consequences of Rab12 knockout in mouse brain and other organs.

### Rab12 overexpression enhances LRRK2 activity

Since loss of Rab12 decreased phosphoRab10 levels, we reasoned that increasing Rab12 should increase phosphoRab10 levels. Indeed, overexpression of GFP-Rab12 in A549 cells led to an almost ten-fold increase in phosphoRab10 levels without changing the levels of LRRK2, PPM1H phosphatase (Berndsen et al., 2019) or total Rab10 (Fig. 3A,B). The ability of Rab12 to activate LRRK2 was specific for that GTPase in that exogenous expression of GFP-tagged Rab8A, Rab10 or Rab29 failed to show the same high level of phosphoRab10 increase—Rab29 yielded about a five-fold enhancement while Rab12 was almost twice as effective in HEK293T cells (Fig. 3C,D). Wild type, R1441C and G2019S LRRK2 expressing cells showed increased phosphorylation upon Rab12 expression (Fig. 3E,F). It is important to note that Rab12 is a much more abundant Rab in most tissues than Rab29—for example, A549 cells contain ∼134000 Rab12 molecules and 25000 Rab29 molecules per cell. This compares with 5000 copies of LRRK2 and 2.5 million copies of Rab10 (https://copica.proteo.info/#/home). Nevertheless, activation was tested at comparable levels of each Rab protein as monitored using anti- GFP-antibodies (Fig. 3C).

**Figure 3.**
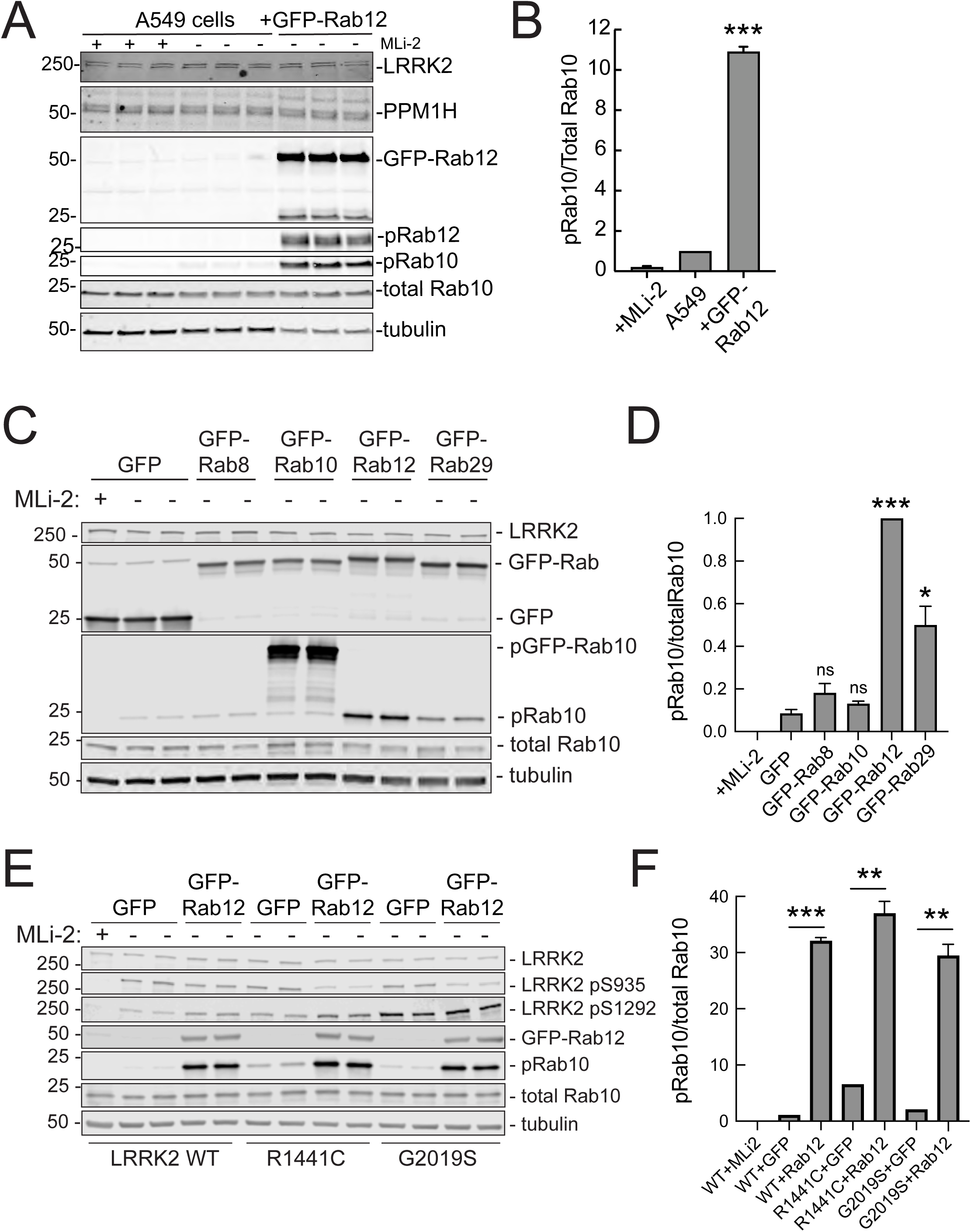
Exogenous Rab12 expression increases phosphoRab10 levels in A549 cells. **(A)** Immunoblot analyses of A549 cells stably overexpressing GFP-Rab12; +/- MLi-2 (200nM for 2h) as indicated. **(B)** Quantitation of the fraction of phosphorylated Rab10 from immunoblots as in **(A)** normalized to total Rab10 levels; error bars indicate SEM from two experiments (***P=0.0003 by Student’s T test). **(C)** Immunoblot analysis of 293T cells transfected with LRRK2 R1441C and GFP, GFP-Rab8, GFP-Rab10, GFP-Rab12, or GFP-Rab29 for 36 hours; +/- MLi2 (200 nM for 2h) as indicated. **(D)** Quantitation of the fraction of phosphorylated Rab10 from immunoblots as in **(C)** normalized to respective total Rab10 levels. Error bars indicate SEM from two independent experiments; ***P = 0.0004 for GFP and GFP-Rab12, *P=0.04 for GFP and GFP-Rab29 with Student’s T test. **(E)** Immunoblot analysis of 293T cells transfected with LRRK2 WT, R1441C or G2019S and GFP or GFP-Rab12 for 36h, +/- MLi2 (200 nM for 2h) as indicated. **(F)** Quantitation of phosphorylated Rab10 from immunoblots as in **(E)** normalized to respective total Rab10 levels. Error bars indicate SEM from two independent experiments; ***P=0.0004 for LRRK2 WT GFP and GFP-Rab12, **P=0.005 for LRRK2 R1441C GFP and GFP-Rab12, **P=0.005 for G2019S GFP and GFP-Rab12 by Student’s T test.

Similar results were obtained using immunofluorescence microscopy to assay phosphoRab10 abundance (Fig. 4). The phoshoRab10 generated was present on perinuclear membrane compartments (Fig. 4A) as seen previously by many groups (cf. Dhekne et al., 2019; 2021; Ordóñez et al., 2019). PhosphoRab10 staining disappeared in cells expressing PPM1H but not in cells expressing the catalytically inactive H153D PPM1H (Fig. 4A,B). These data were confirmed by immunoblot (Fig. 4C,D) and suggest that Rab12 is activating LRRK2 along the same pathway of protein phosphorylation studied previously to date.

**Figure 4.**
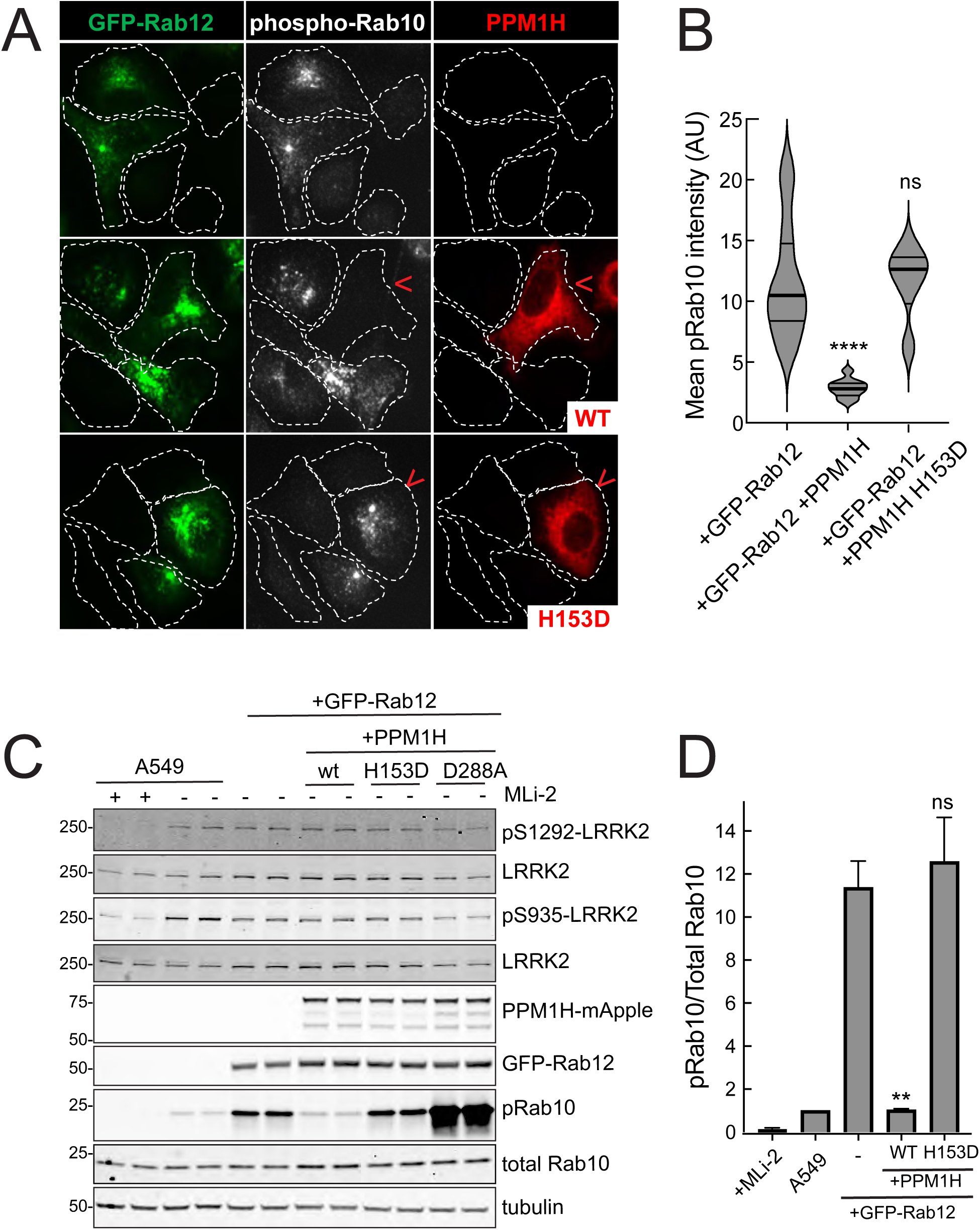
PPM1H phosphatase counters phosphoRab10 generated upon Rab12 activation. **(A)** A549 cells stably expressing GFP-Rab12 and PPM1H-mApple (wildtype and H153D catalytically inactive mutant) were co-cultured with parental wild type A549 cells on coverslips. PhosphoRab10 was detected by immunofluorescence using rabbit anti-phosphoRab10. Red arrowheads indicate a cell with both GFP-Rab12 and wtPPM1H-mApple or PPM1H H153D. **(B)** Quantitation of mean phosphoRab10 fluorescence intensity per cell (Arbitrary units, AU) is shown in the violin plot. Error bars indicate SEM from two independent experiments. At least 10 cells per condition were counted.****P <0.0001 for GFP- Rab12 and GFP-Rab12+wt PPM1H, ^ns^P=0.9944 for GFP-Rab12 and GFP-Rab12 + [H153D]PPM1H with Student’s T-test. **(C)** Immunoblot analysis of parental A549 cells or A549 cells stably expressing GFP-Rab12 together with either wt PPM1H, [H153D] PPM1H or [D288A] PPM1H; +/- MLi2 (200 nM for 2h) as indicated. **(D)** Quantitation of the fraction of phosphorylated Rab10 from immunoblots as in **A** normalized to respective total Rab10 levels. Error bars indicate SEM from two independent experiments; **P=0.007 for, GFP-Rab12 and GFP-Rab12+wt PPM1H, ^ns^P=0.5510 for GFP-Rab12 and GFP-Rab12+[H153D] PPM1H by Student’s T-test.

### Requirements for Rab12 activation of the LRRK2 pathway

It was possible that Rab12 activated a kinase other than LRRK2 to increase Rab10 phosphorylation. This appears not to be the case as GFP-Rab12 expression enhancement of phosphoRab10 levels was not seen in A549 cells lacking LRRK2 expression (Fig. 5A,B). It was possible that exogenous GFP- Rab12 inhibited overall Rab phosphatase activity, leading to an apparent increase in phosphoRab10 levels. This was also ruled out, as cells lacking PPM1H displayed full Rab12-induced enhancement of phosphoRab10 levels (Fig. 5C,D), about five-fold with or without PPM1H.

**Figure 5.**
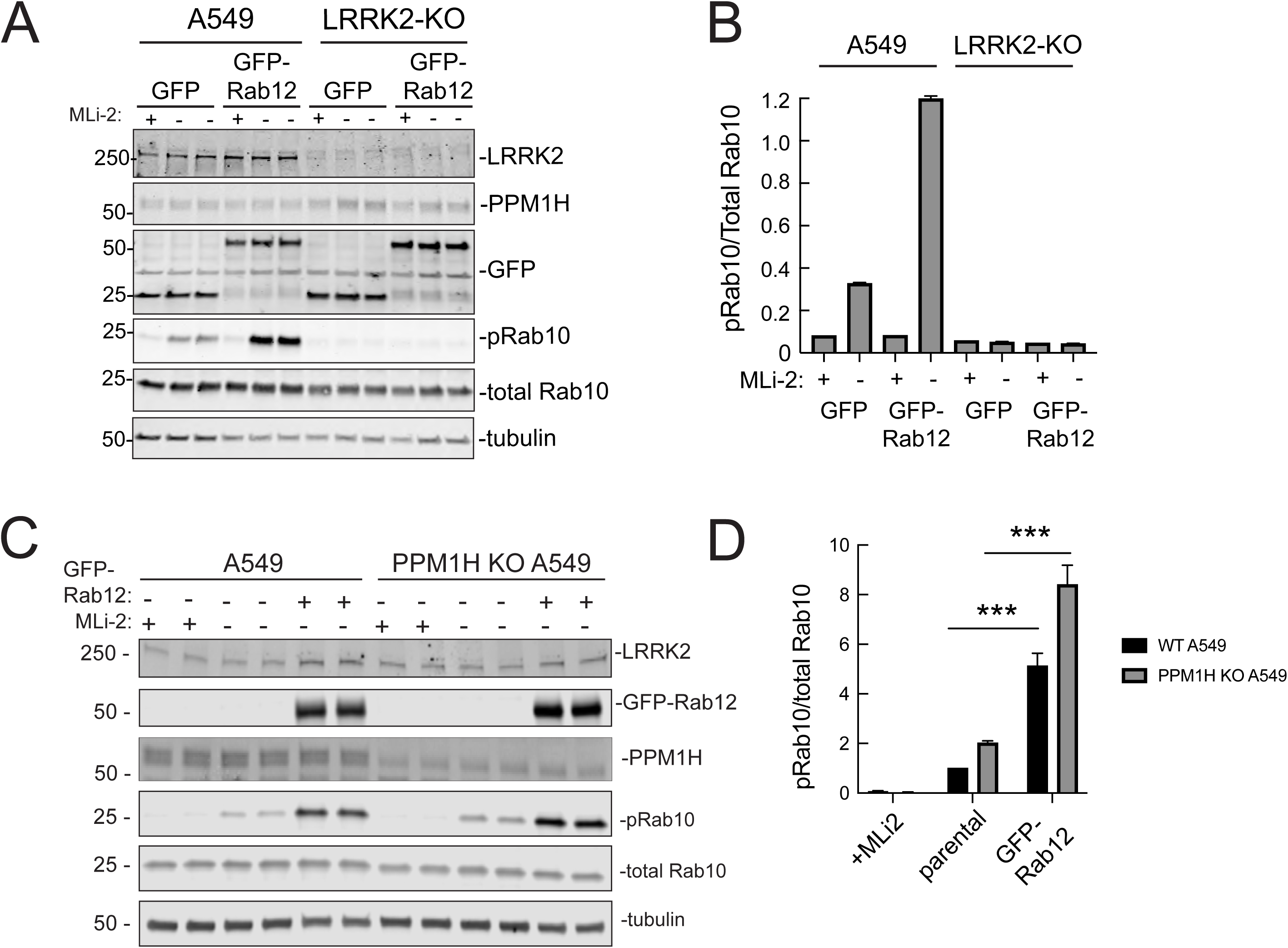
Roles of LRRK2 and PPM1H in Rab12 activation of LRRK2. **(A)** Immunoblot analysis of WT and LRRK2 KO A549 cells stably expressing GFP or GFP-Rab12; +/- MLi-2 (200 nM for 2h) as indicated. **(B)** Quantitation of the fraction of phosphorylated Rab10 from immunoblots as in **A** normalized to respective total Rab10 levels. **(C)** Immunoblot analysis of WT and PPM1H KO A549 parental cells or cells stably expressing GFP-Rab12; +/- MLi-2 (200 nM for 2h) as indicated. **(D)** Quantitation of the fraction of phosphorylated Rab10 from immunoblots as in C normalized to respective total Rab10 levels, normalized to WT parental. Error bars indicate SEM from four independent experiments; ***P=0.0002 for both WT and PPM1H KO parental and GFP-Rab12 by Student’s T test.

### Rab12 activation requires a novel Rab binding site in the LRRK2 Armadillo domain

Previous work has identified specific residues within LRRK2 Armadillo domain that enable LRRK2 to be recruited to the Golgi by exogenously overexpressed Rab29; these residues support direct Rab29 binding (McGrath et al., 2021; Vides et al., 2022; Zhu et al., 2022). In particular, R361, R399, and K439 contribute to a Rab binding “Site #1” that supports binding to purified Rab29 (K_D_=1.6µM; Vides et al., 2022; see Fig. 9 below). Rab8A binds this LRRK2 350-550 region with a similar affinity (2.3µM) but Rab10 binds less well (5.1µM) (Vides et al., 2022); a second site at LRRK2’s N-terminus (K17/K18) mediates interaction with phosphorylated Rab8A and Rab10 protein.

AlphaFold (Jumper et al., 2021) in conjunction with Colabfold in ChimeraX (Mirdita et al., 2022; Pettersen et al., 2004) revealed a third Rab binding site (Site #3) when the full-length Armadillo domain was modeled together with Rab12 (Fig. 6A; see Fig. 9 below). Mutagenesis across this putative binding interface yielded full length LRRK2 proteins with decreased overall activity as monitored by phosphoRab10 levels in HEK293T cells expressing the mutant proteins (Fig. 6B). Note that these cells rely only on endogenous Rab12 protein for possible LRRK2 activation. Thus, mutation of E240 and S244 had the greatest impact on LRRK2 activity; remarkably, mutation of F283 to A increased kinase activity two-fold. These data demonstrate that Site #3 sequences are important for overall LRRK2 activity.

**Figure 6.**
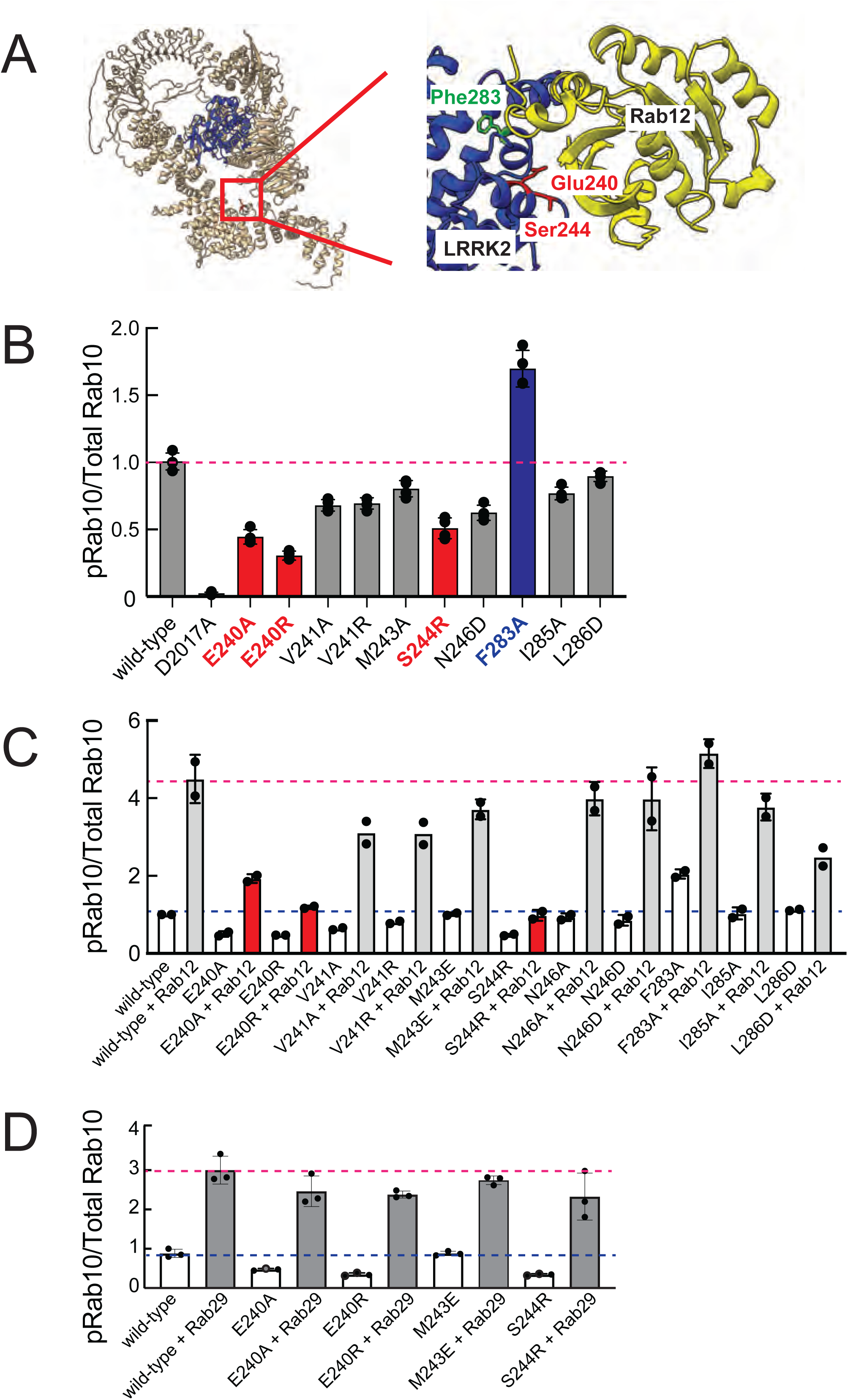
Rab binding Site 3 is needed for Rab12- but not Rab29-mediated LRRK2 activation. **(A)** AlphaFold model highlighting the residues at the interface between LRRK2 and Rab12 **(B)** Immunoblot analysis of 293 cells transfected with the indicated LRRK2 “Site#3” mutants. The quantitation of the fraction of phosphorylated Rab10 from immunoblots as in Figure 6-Figure Supplement 1 normalized to respective total Rab10 levels is shown. **(C)** Immunoblot analysis of 293 cells co-transfected with the indicated LRRK2 “Site#3” mutants and HA or HA-Rab12. The quantitation of the fraction of phosphorylated Rab10 from immunoblots as in Figure 6-Figure Supplement 1 normalized to respective total Rab10 levels is shown. **(D)** Immunoblot analysis of 293 cells co-transfected with the indicated LRRK2 “Site#3” mutants and HA or HA-Rab29. The quantitation of the fraction of phosphorylated Rab10 from immunoblots as in Figure 6-Figure Supplement 1 normalized to respective total Rab10 levels is shown.

Mutation of LRRK2 E240R and S244R predicted to be important for Rab12 binding blocked the ability of exogenous Rab12 to enhance phosphoRab10 levels (Fig. 6C and Fig. 6–Figure Supplement 1).

Moreover, F283A LRRK2 had twofold higher basal activity but was not activated by exogenous Rab12 significantly more than wild type LRRK2 protein. These data strongly suggest that Rab12 activates LRRK2 by binding to Site #3 within the Armadillo domain.

Extensive previous mutagenesis defined Site #1 as being critical for exogenous Rab29-dependent relocalization of LRRK2 to the Golgi complex and apparent activation (Vides et al., 2022). It was therefore important to assess whether Rab29’s ability to increase phosphoRab10 levels upon overexpression relied upon Site #3. As expected, exogenous expression of Rab29 increased phosphoRab10 levels (albeit to a lower extent than exogenous Rab12 expression (Fig. 3C,D; Fig. 6D). However, mutation of Site #3 residues critical for Rab12-mediated LRRK2 activation (E240 and S244) had no effect on the ability of Rab29 to activate LRRK2 kinase. These experiments indicate that Rab29 interacts preferentially with Site #1 and demonstrate the Rab12 selectivity of Site #3 for LRRK2 activation.

These experiments strongly suggest that Rab29 and Rab12 activate LRRK2 by two different routes: Rab29 via LRRK2 Site #1 and Rab12 via Site #3. We validated Rab12 direct binding to Site # 3 using purified Rab12 and Armadillo domain proteins mutated at either Site #1 (K439E) or Site #3 (E240R). As shown in Figure 7, Rab12 bound as well to the wild type Armadillo domain (Fig. 7A, 1.4µM) as to an Armadillo domain construct bearing a Site #1 mutation (Fig. 7B, 1.6µM) as determined by microscale thermophoresis. In contrast, the Site #3 E240R mutation abolished the interaction, yielding a K_D_ of >40µM (Fig. 7C). These data show that Rab12 binds tightly and directly to Site #3 in vitro and does not appear to interact significantly with Site #1.

**Figure 7.**
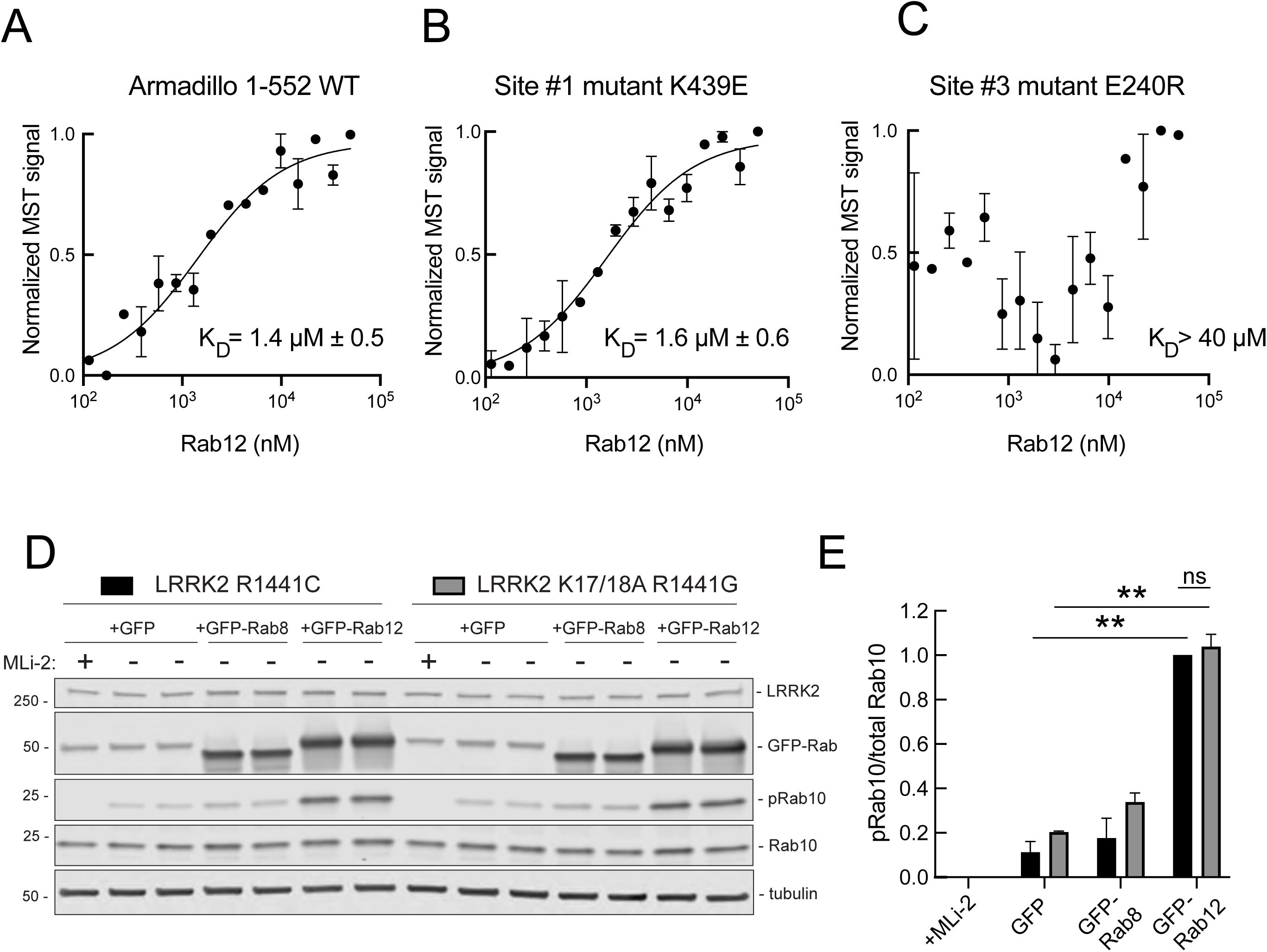
Rab12 binds directly to Site 3 and Site 2 is dispensable for Rab12-mediated LRRK2 activation. **(A, B, C)** Microscale thermophoresis of Rab12 binding to fluorescently labeled LRRK2 Armadillo domain (residues 1–552) wild type (A) or bearing the indicated mutations at Site #1 **(B)** or Site #3 **(C)**. Purified Rab12 was serially diluted and then NHS-RED-labeled-LRRK2 Armadillo (final concentration 100 nM) was added. Graphs show mean and SEM from two independent measurements, each the average of two replicate runs.(**D)** Immunoblot analysis of 293T cells transfected with LRRK2 R1441C or K17/18A R1441G and GFP, GFP-Rab8, or GFP-Rab12 for 36 hours; +/- MLi2 (200 nM for 2h). **(E)** Quantitation of the fraction of phosphorylated Rab10 from immunoblots as in **(F)** normalized to respective total Rab10 levels, normalized to LRRK2 R1441C + GFP-Rab12. Error bars indicate SEM from two independent experiments; **P=0.003 for LRRK2 R1441C GFP and GFP-Rab12, **P=0.0044 for LRRK2 K17/18A R1441G GFP and GFP-Rab12, ns=0.6 by Student’s T-test.

### PhosphoRab binding is distinct from the Rab12 pathway of LRRK2 activation

We showed previously that phosphoRab binding to Rab binding Site #2 is critical for cooperative LRRK2 membrane recruitment and apparent activation (Vides et al., 2022). Thus, it was important to investigate whether Rab12 acts via this feed-forward process: it was possible that phosphoRab12 was driving activation rather than non-phosphorylated Rab12. If true, such activation would be predicted to rely on LRRK2 Lys17 and Lys18. As shown in Figure 7D,E, mutation of Lys17 and 18 had no effect on the ability of Rab12 to increase phosphoRab10 levels in HEK293T cells co-expressing exogenous LRRK2 and GFP-Rab proteins. Once again, GFP-Rab12 activation was dramatic and K17/K18 containing-LRRK2 was activated to the same overall level as the K17A/K18A mutant LRRK2 protein.

Finally, an alternative way to investigate this question would be to use a non-phosphorylatable Rab12 protein. Unfortunately, this approach cannot be taken, as non-phosphorylatable Rabs 8A and 10 do not behave like their wild type counterparts: Rab8A- and Rab10-TA are mis-localized and cannot rescue corresponding ciliary phenotypes; moreover, Rab10-TA cannot be prenylated in vitro, indicative of a nucleotide binding issue (Dhekne et al., 2018). Results from experiments using such mutants must be interpreted with great care and deeper analysis.

### Rab12 drives LRRK2 activation upon lysosomal or ionophore-triggered stress

As mentioned earlier, under conditions of lysosomal damage, LRRK2 is recruited to lysosomes and participates in the repair of damaged endomembranes (cf. Eguchi et al., 2018; Herbst et al., 2020; Bonet-Ponce, 2020). Such stress greatly increases LRRK2 kinase activity (cf. Kalogeropulou et al., 2020). Figure 8 shows that Rab12 is required for the increase in LRRK2 activity upon lysosomal damage triggered by LLOME addition (Fig. 8A-C) or especially upon treatment of cells with Nigericin that also causes mitochondrial stress and is a potent activator of the NLRP3 inflammasome (Fig. 8D-F). These findings point to the importance of Rab12 in regulating LRRK2 activity in lysosome repair.

**Figure 8.**
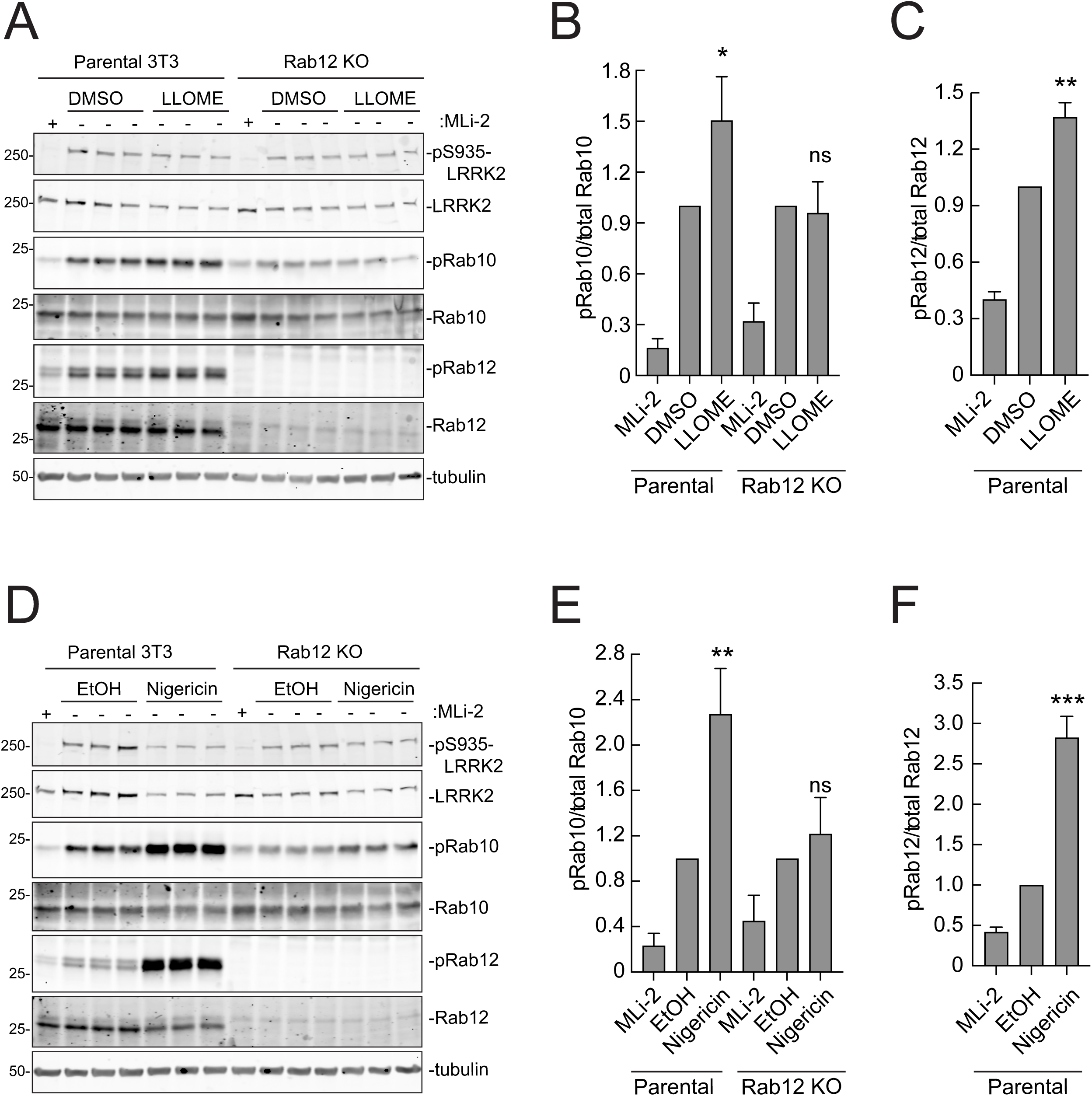
Rab12 is needed for LRRK2 activation by LLOME and Nigericin. **(A)** Immunoblot analyses of WT and Rab12 KO 3T3 cells treated with 1mM LLOME for 2h, +/- MLi-2 (200nM for 2h) as indicated. **(B)** Quantitation of phosphorylated Rab10 from immunoblots as in **A** normalized to total Rab10; Error bars indicate SEM from three experiments (*P=0.0282, ^ns^P=0.7189 by Student’s T test). **(C)** Quantitation of phosphorylated Rab12 from immunoblots as in **A** normalized to total Rab12 levels; Error bars indicate SEM from three experiments (**P=0.0011 by Student’s T test). **(D)** Immunoblot analyses of WT and Rab12 KO 3T3 cells treated with 2µM LLOME for 2 h, +/- MLi-2 (200nM for 2h) as indicated. **(E)** Quantitation of phosphorylated Rab10 from immunoblots as in **D** normalized to total Rab10 levels; Error bars indicate SEM from three experiments (**P=0.0054, ^ns^P=0.3100 by Student’s T test). **(F)** Quantitation of the fraction of phosphorylated Rab12 from immunoblots as in **D** normalized to total Rab12 levels; Error bars indicate SEM from three experiments (***P=0.0003 by Student’s T test).

## Discussion

Using an unbiased, genome-wide screen, we have discovered an important and unanticipated role for the understudied Rab12 GTPase in LRRK2 kinase regulation. Loss of Rab12 from NIH3T3 and MEF cells (and possibly also mouse lung tissue) significantly decreased phosphoRab10 levels, and Rab12 overexpression increased phosphoRab10 levels. The phosphoRab10 increase was LRRK2 dependent, Rab12 specific, and seen with both wild type and pathogenic mutant LRRK2 proteins. PhosphoRab10 showed the same subcellular localization seen in prior work with cells expressing hyperactive LRRK2 proteins and was sensitive to the Rab-specific, PPM1H phosphatase, consistent with Rab12 activation being part of the normal LRRK2 phosphorylation pathway. Site directed mutagenesis in conjunction with computational modeling revealed a new Rab binding site (#3) within the LRRK2 Armadillo domain that is needed for Rab12 binding and activation and is not engaged by Rab29 to trigger apparent kinase activation.

Figure 9 summarizes our current knowledge of Rab GTPase Armadillo domain interactions. Rab29 and its relatives, Rab32 and Rab38, can bind to Site #1 that includes LRRK2 R361, R399, and K439 residues (McGrath et al., 2021; Vides et al., 2022; Zhu et al., 2022); Rab8A is also able to bind at that location (Vides et al., 2022). PhosphoRab8A and phosphoRab10 interact with comparable high affinity with LRRK2 K17/18 at Site #2. This study reveals a third interaction interface E240, S244) on the opposite face of the Armadillo domain relative to Site #1 that engages Rab12 GTPase. The cryoEM structures of full length LRRK2 (Myasnikov et al., 2021) or LRRK2 in the presence of Rab29 (Zhu et al., 2022) both show an extended and flexible Armadillo domain that extends away from the kinase center and would be available for Rab GTPase engagement.

**Figure 9.**
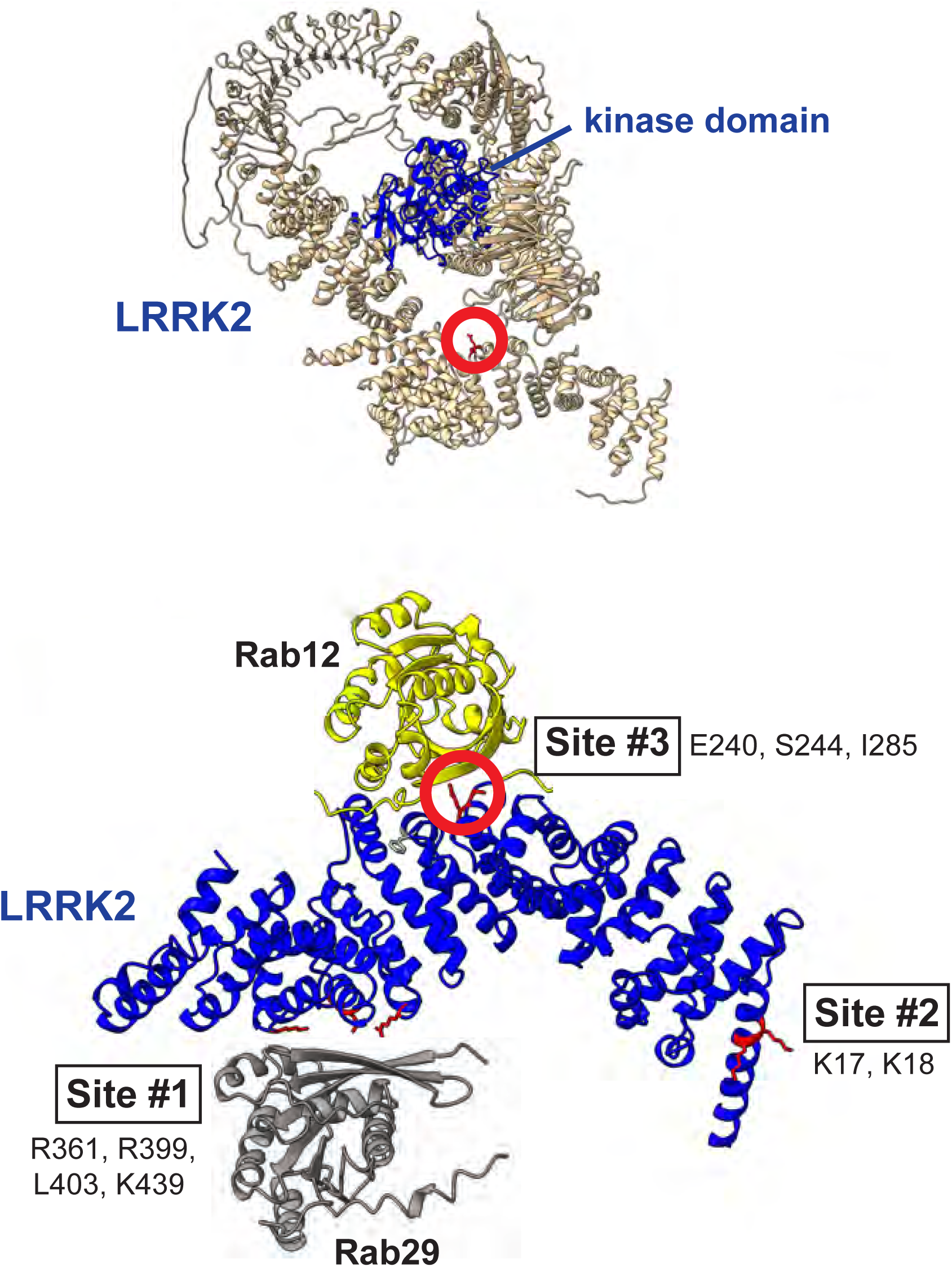
Model for Rab interactions with LRRK2 Armadillo domain. Shown at the bottom is the AlphaFold model for LRRK2 Armadillo (blue) interaction with Rab12 (yellow) and Rab29 (gray). The Rab12 was docked onto Armadillo using Colabfold in ChimeraX; Rab29 was positioned manually. Site 1 binds Rab29; Site 2 binds phosphorylated Rabs (Vides et al., 2022) and Site 3 binds Rab12. The key residue for Rab12 binding is circled in red in both the lower Armadillo domain model and in the corresponding location within the full length LRRK2 Alphafold model shown above.

What are the roles of these multiple Rab binding sites? Site #1 can interact with overexpressed Rab29 protein and bring the mostly cytosolic LRRK2 kinase to the surface of the Golgi complex, which will lead to apparent activation. With regard to membrane anchoring, since loss of Rab29 has no detectable consequence for Rab phosphorylation (Kalogeropulou et al., 2020), it seems likely that Site #1 can also be occupied by the ubiquitous and more abundant Rab8A GTPase. Site #2 that binds to phosphoRabs will also contribute to the membrane anchoring of LRRK2 kinase (Vides et al., 2022). Site #3 faces the kinase domain in the AlphaFold model of a putative active LRRK2 protein (Figure 6; Fig. 9) and we propose that Rab12 binding to Site #3 holds open the kinase for full catalytic activity. Given that Rab12’s Ser106 phosphorylation site faces the Armadillo domain as part of this protein binding interaction, LRRK2 contains at least one additional, yet to be discovered, substrate binding site that positions the Rab phosphorylation site in the correct orientation for LRRK2 kinase phospho-addition.

Rabs 8A, 10 and 12 do not co-localize in cells yet they can all interact with LRRK2. One possibility is that LRRK2 binds one Rab in each compartment, independently. If Rab8 recruits LRRK2, Rab8 and phosphoRab8 will both cooperate to hold LRRK2 on a Rab8-enriched membrane surface. How would Rab12 come in? It is important to keep in mind the fact that in an A549 cell with 134,000 Rab12 molecules and ∼ 1 million Rab8A proteins, the 5,000 LRRK2 molecules (https://copica.proteo.info/#/home) may find a subcompartment that contains both Rab8A or 10 and Rab12, despite different primary localizations for the bulk of these Rab proteins. It is also possible that LRRK2 recruited by a Rab to one membrane compartment can phosphorylate a Rab on an adjacent membrane compartment. Future relocalization experiments such as those that anchor LRRK2 on specific subcellular compartments (cf. Gomez et al., 2019; Kluss et al., 2022) may shed important light on this interesting question.

Beyond activating LRRK2, little else is known about Rab12 GTPase function. In mast cells, Rab12 promotes microtubule-dependent retrograde transport of secretory granules (Efergan et al., 2016). GFP-Rab12 co-localizes with transferrin receptors and the PAT4 amino acid transporter and depletion of Rab12 increases the levels of both of these proteins, leading Fukuda and colleagues to conclude that it functions in membrane protein delivery from the endocytic recycling compartment to lysosomes (Matsui & Fukuda, 2011; 2013; Matsui et al., 2011). These workers showed further that Rab12 regulates the constitutive degradation of PAT4, indirectly influencing mTORC1 activity by modulating cellular amino acid levels. Later work from McPherson showed that under starvation conditions, the DENND3 Rab12 guanine nucleotide exchange factor is phosphorylated by ULK kinase, enhancing its activity and overall levels of Rab12-GTP (Xu et al., 2015). Future work will investigate the consequences of starvation on Rab12 localization and possible roles in autophagy and ciliogenesis regulation. LRRK2 is recruited to damaged lysosomes such as those seen in cells treated with lysosomotropic agents or the LLOME peptide (Eguchi et al., 2018; Herbst et al., 2020; Bonet-Ponce et al., 2020). As we show here, Rab12 also plays a critical role in activating LRRK2 in that context.

Pathogenic mutations in LRRK2 kinase cause Parkinson’s disease, and LRRK2 kinase inhibitors are currently in clinical trials in the hopes of benefiting patients (cf. Jennings et al., 2022). This work suggests that small molecules that interfere with Rab12 binding to LRRK2 or other means that decrease Rab12 levels may provide additional avenues to target hyperactive LRRK2 kinase.

---

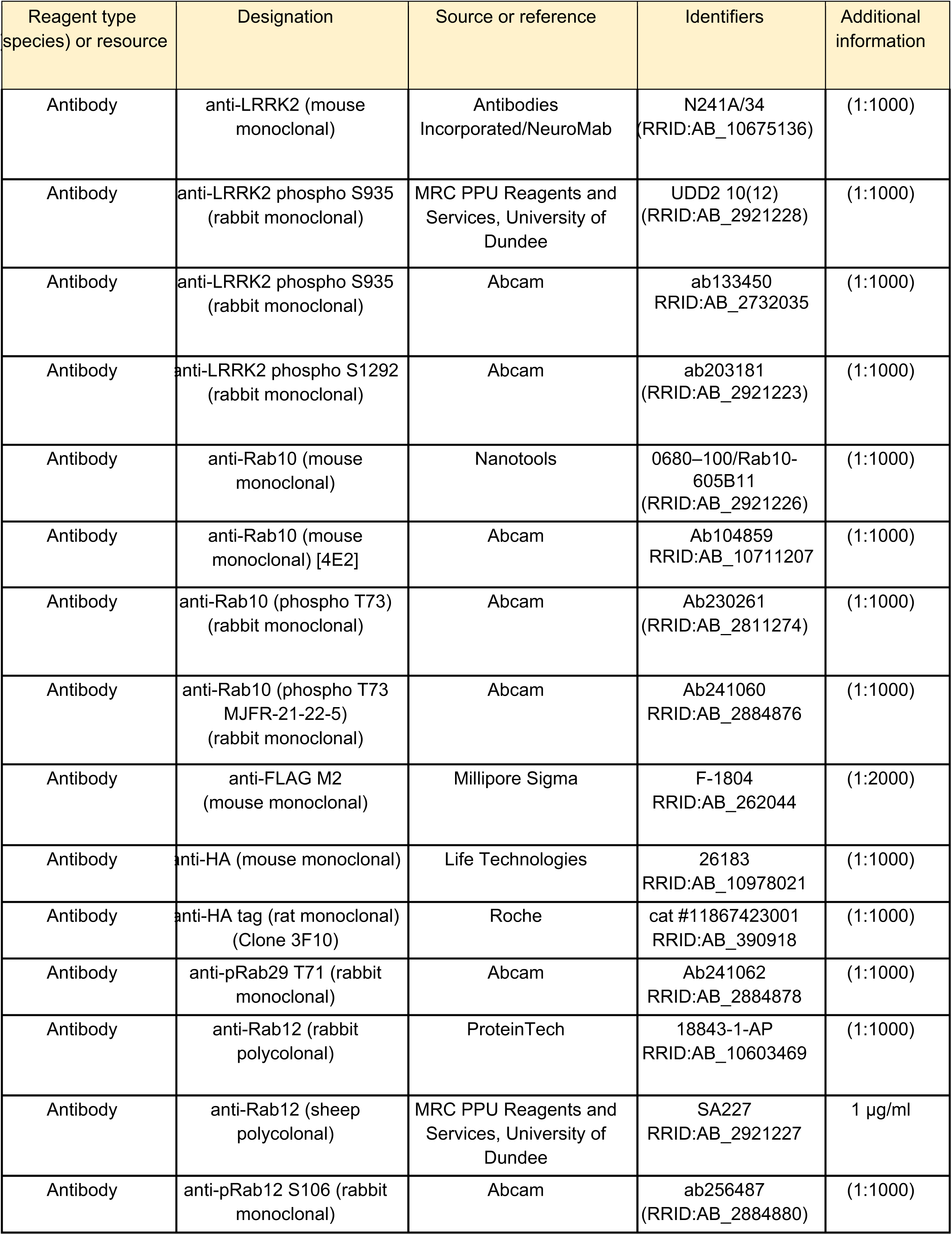
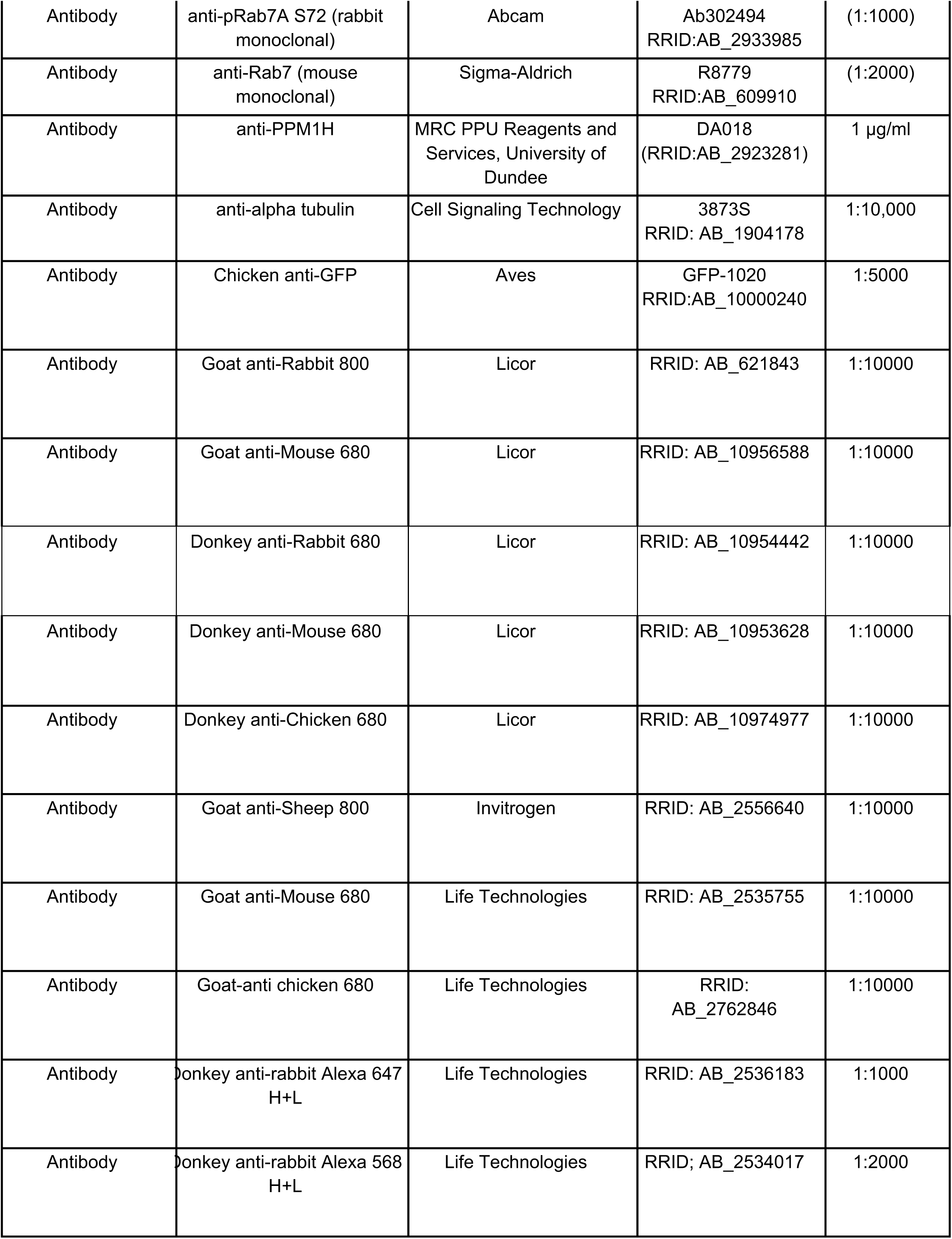
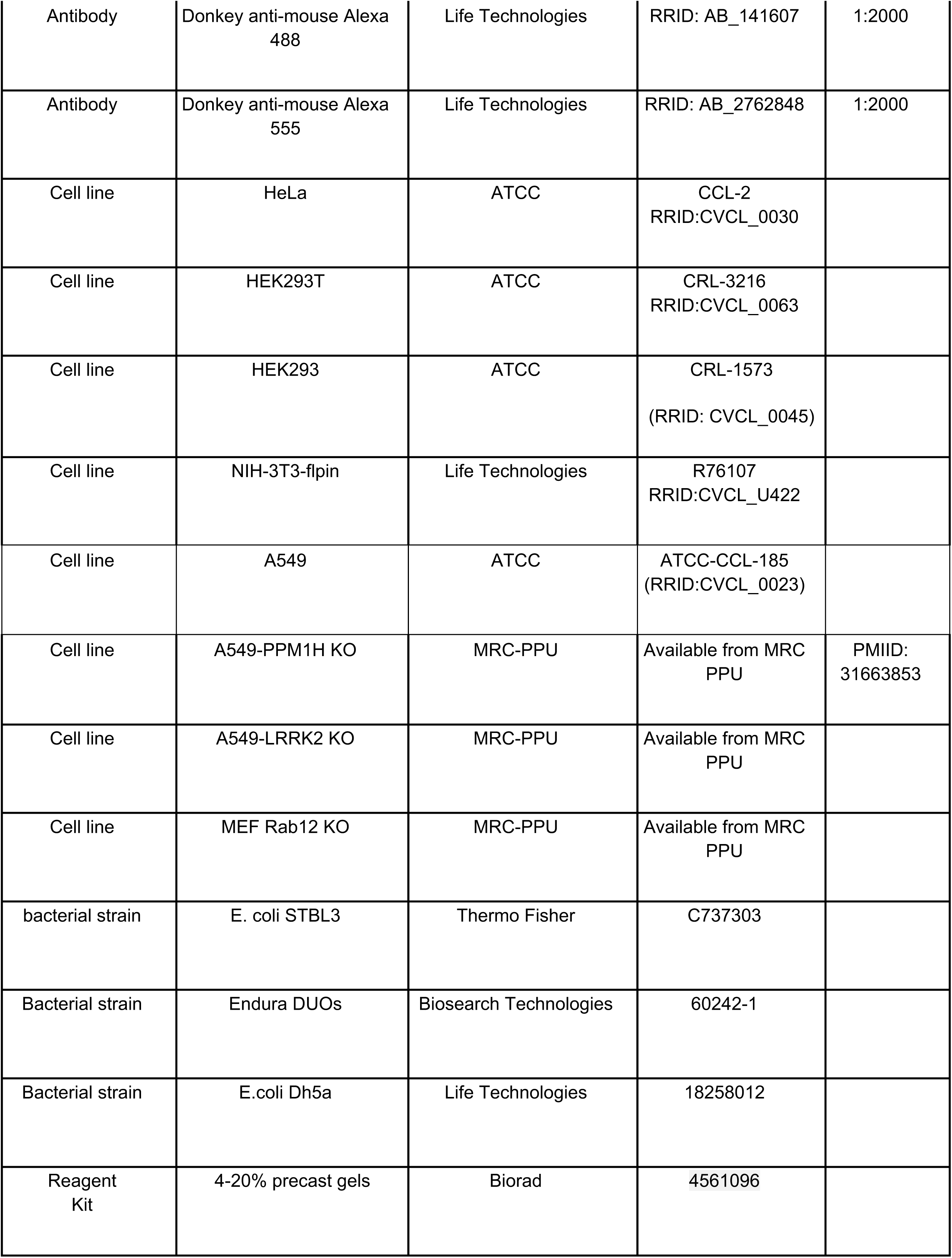
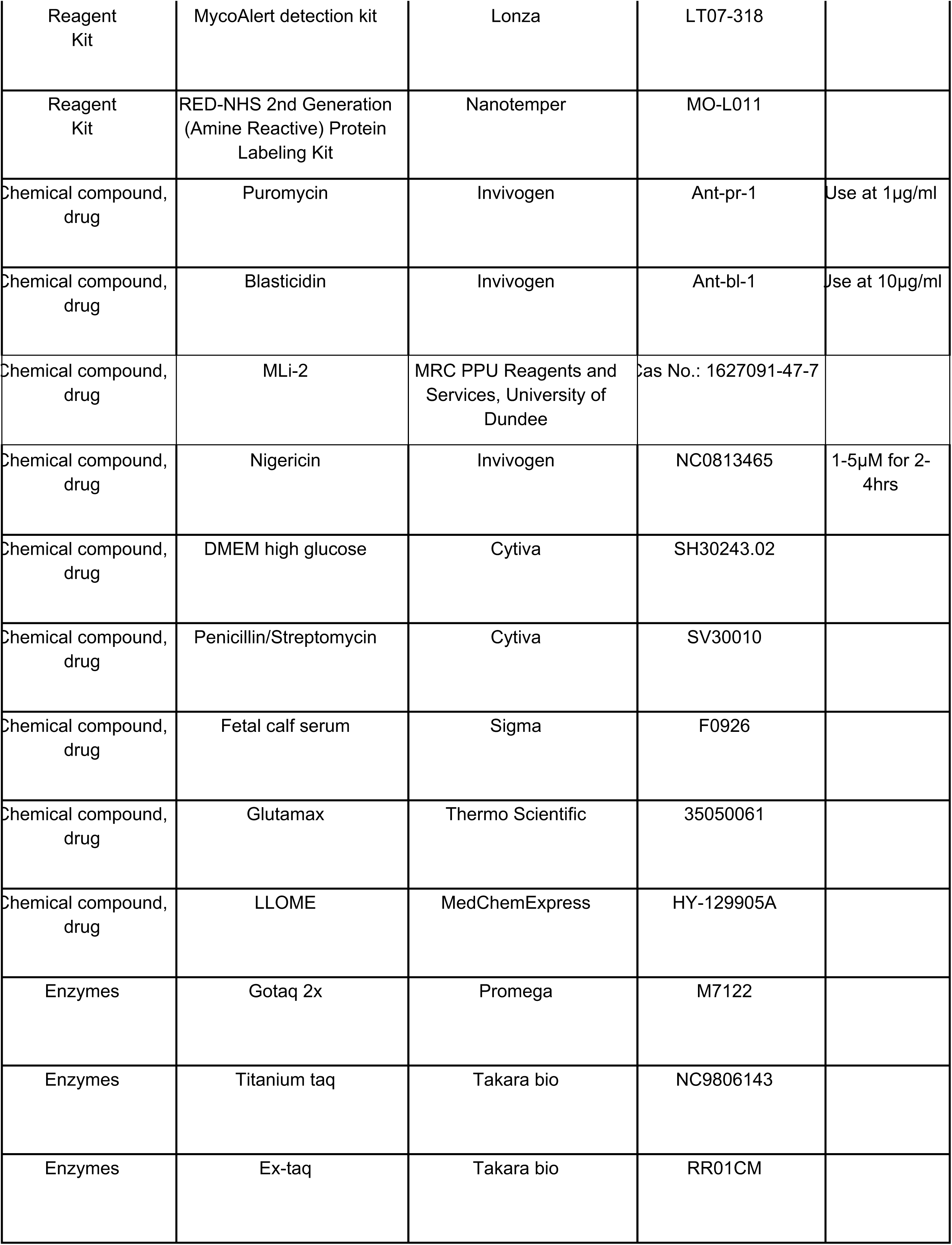
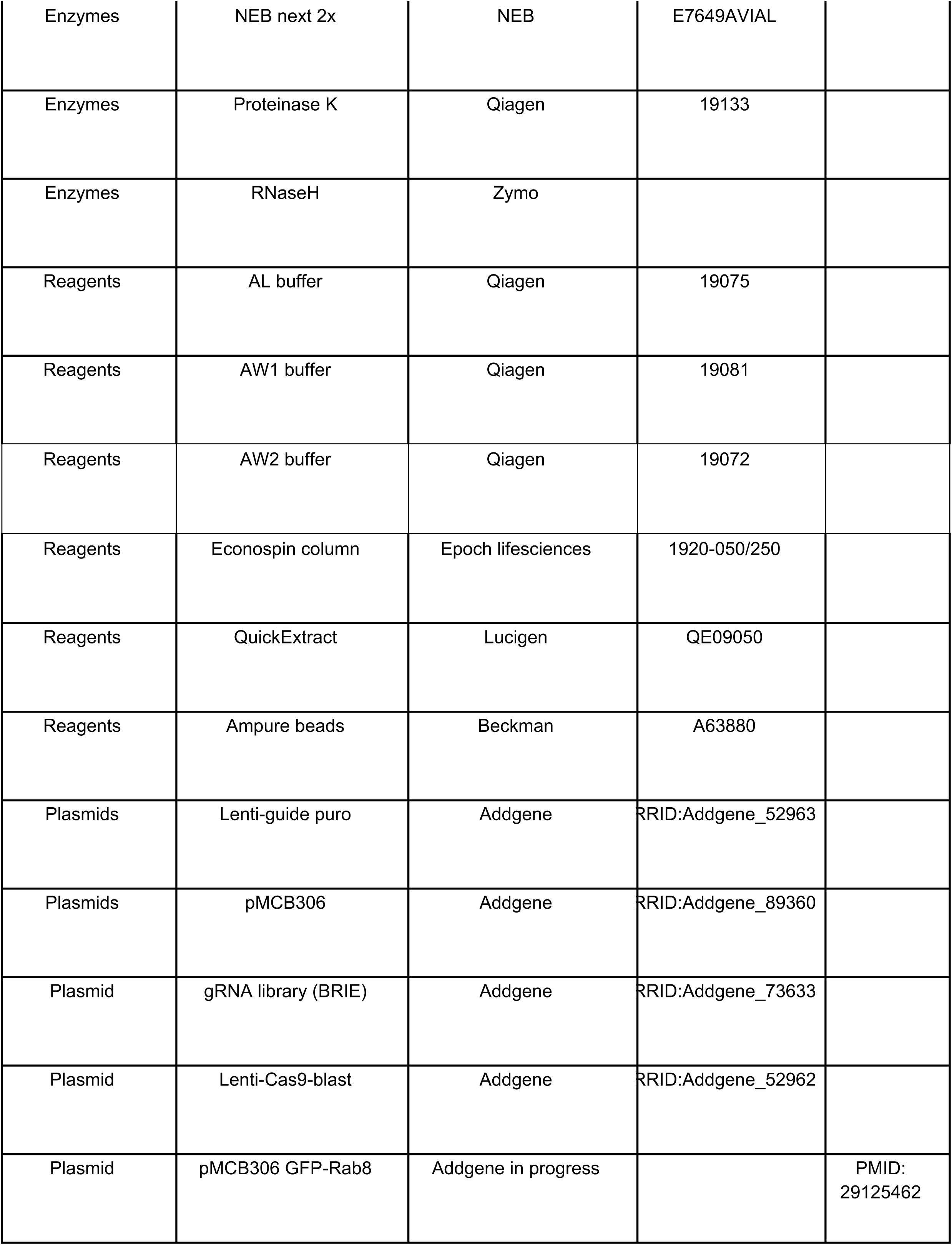
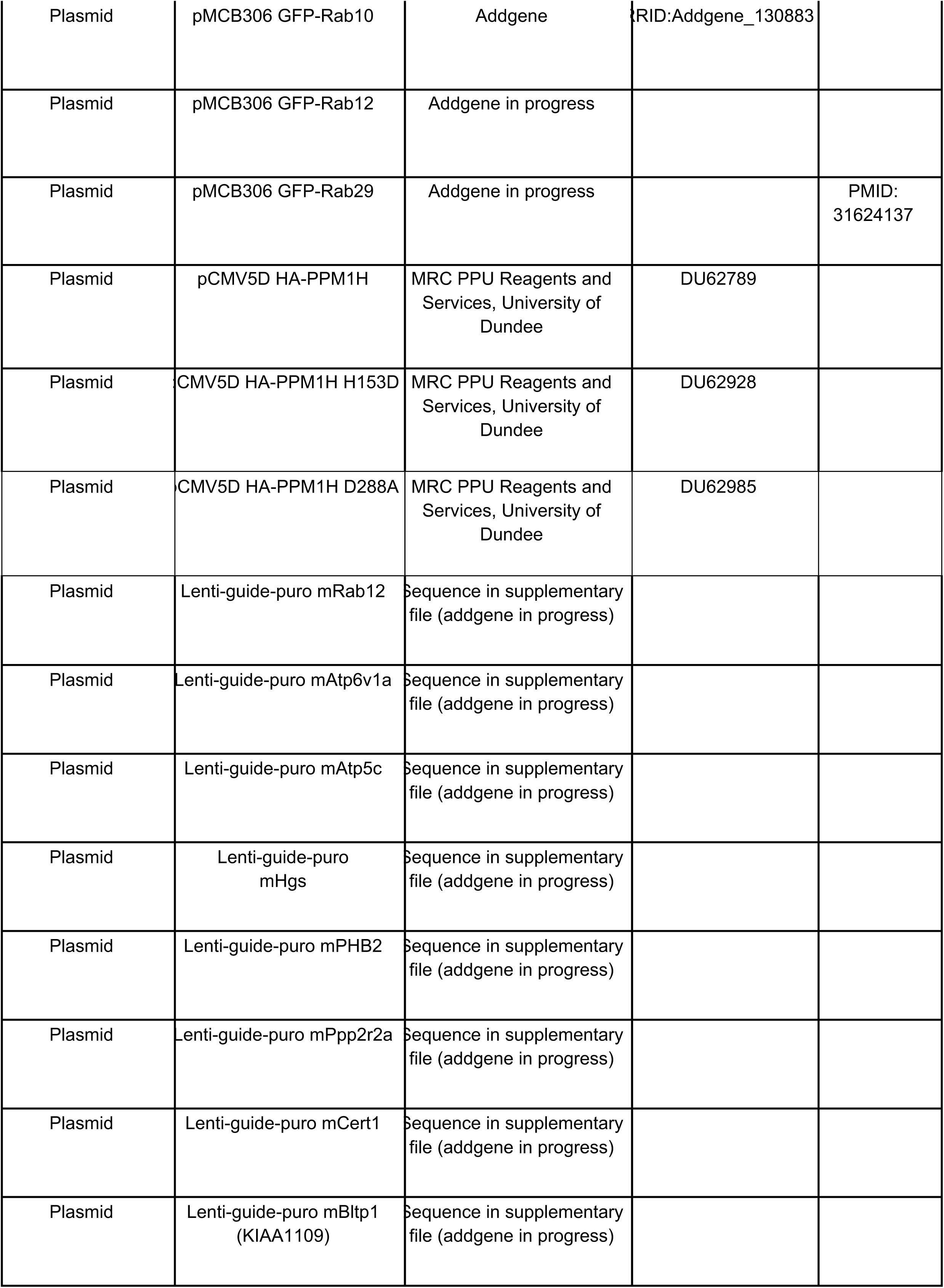
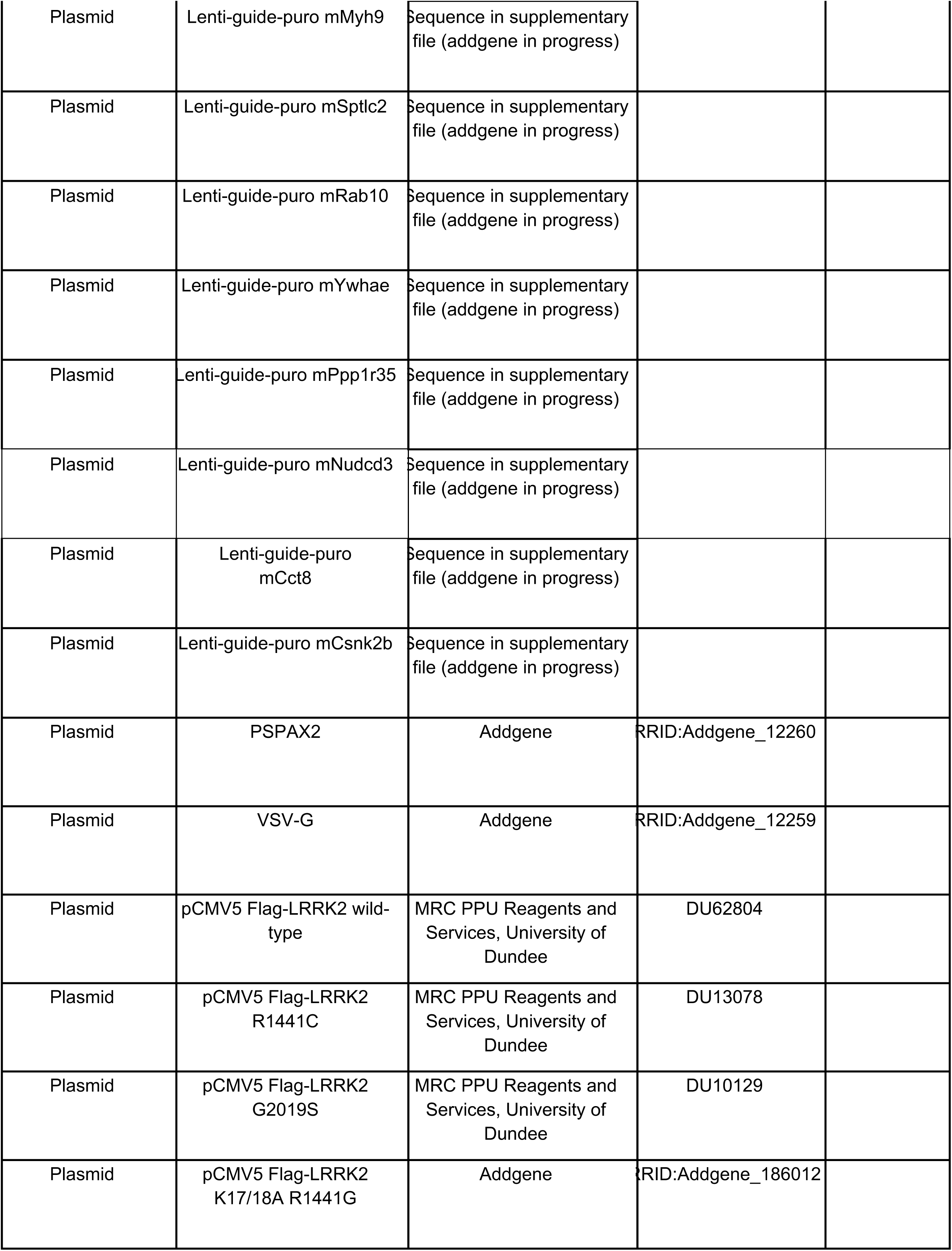
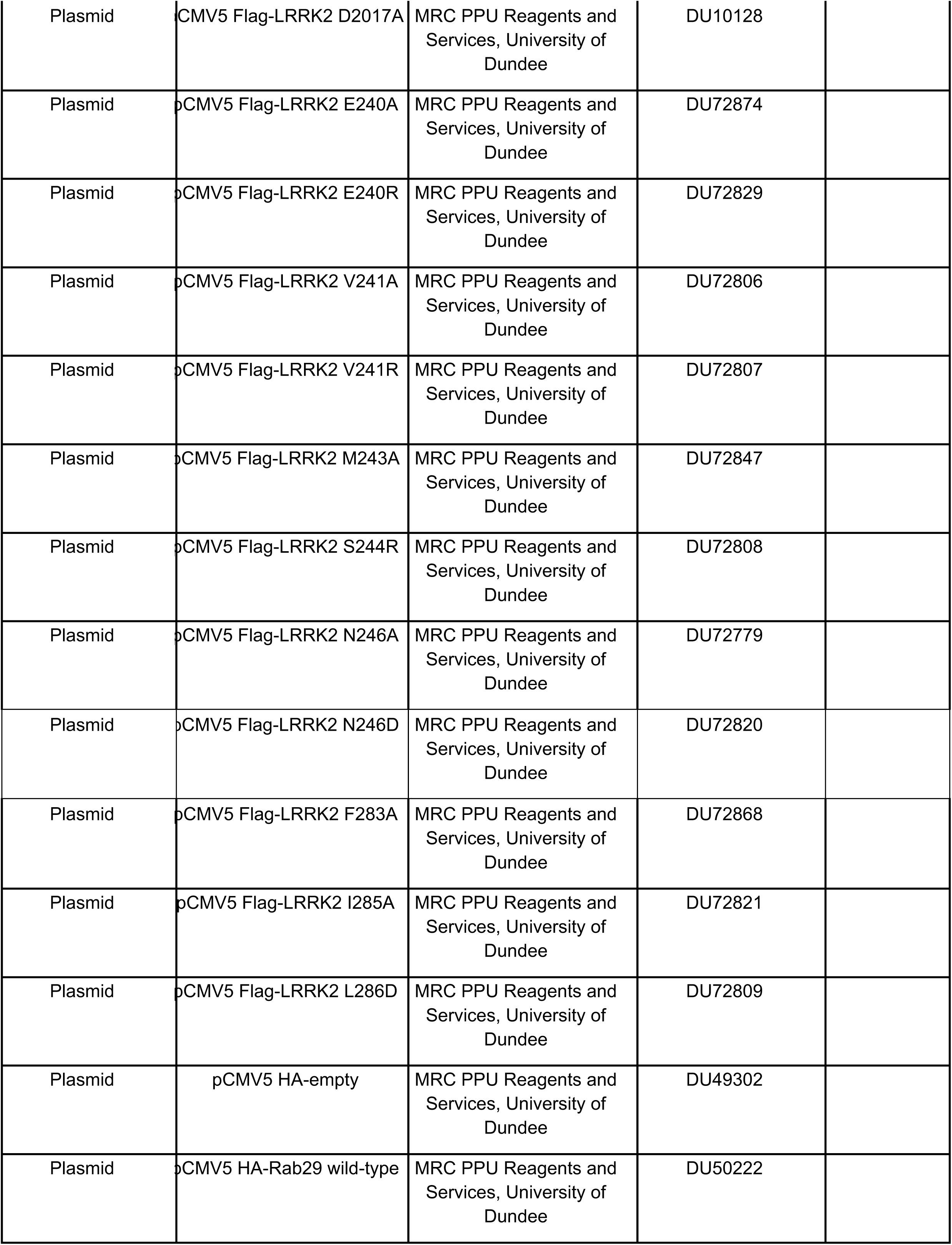
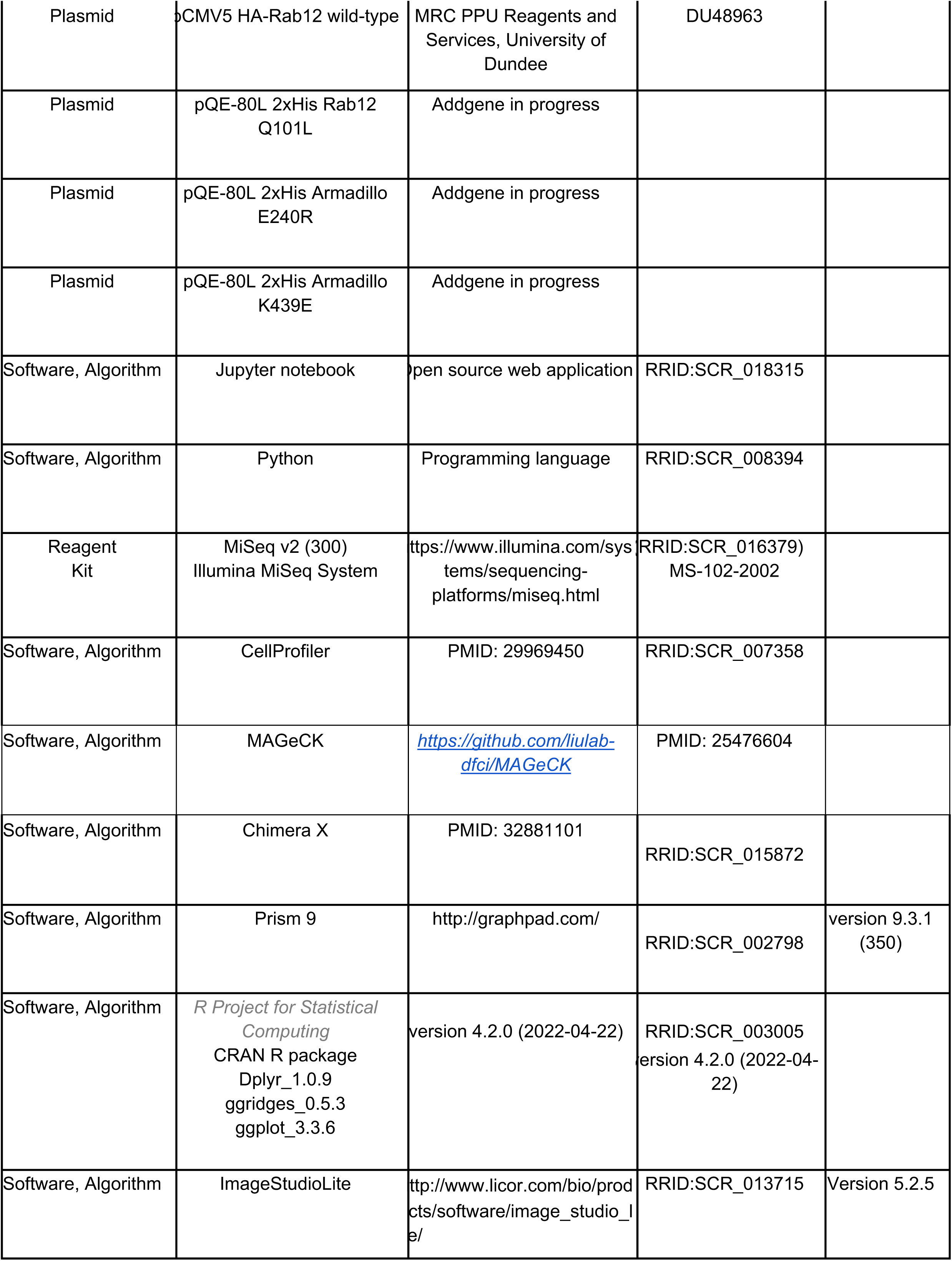
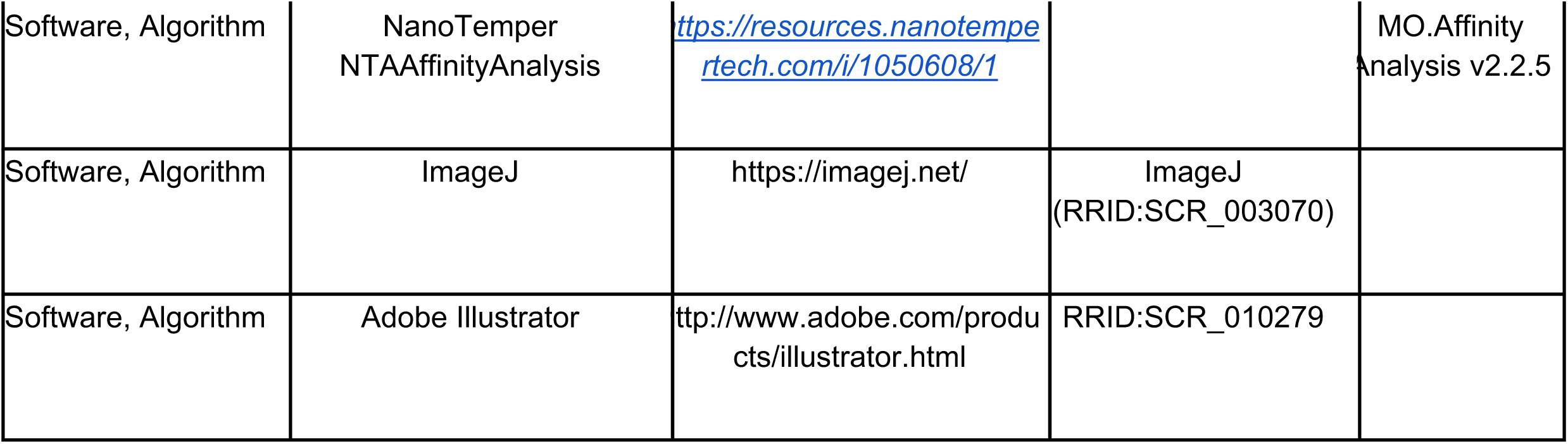
Key resources

## Acknowledgements

This study was funded by the joint efforts of The Michael J. Fox Foundation for Parkinson’s Research (MJFF) (MJFF grant no. 009258 to SRP and DRA and 021132 to SRP) and Aligning Science Across Parkinson’s (ASAP) initiative. MJFF administers the grant (ASAP-000463, SRP and DRA) on behalf of ASAP and itself. CYC was supported by training grant NIH 5 T32 GM007276. Funds were also provided by the Medical Research Council (grant no. MC_UU_00018/1 [DRA]), the pharmaceutical companies supporting the Division of Signal Transduction Therapy Unit Boehringer-Ingelheim, GlaxoSmithKline, Merck KGaA (DRA). For the purpose of open access, the authors have applied a CC-BY public copyright license to the Author Accepted Manuscript version arising from this submission. All primary data associated with each figure has been deposited in a repository and can be found at doi.org/10.5281/zenodo.7633918 and XXXXXXX.

We are especially grateful to Drs. Ganesh Puspati and Rajat Rohatgi for critical guidance in performing the 3T3 cell CRISPR screen, Jacqueline Bendrick and Yohan Auguste for help with Fig. 7A-C, and Dr. Sreeja Nair for help sustaining clones while HSD recovered from COVID. We also thank the excellent technical support of the MRC protein phosphorylation and ubiquitylation unit (PPU) DNA sequencing service (coordinated by Gary Hunter), the MRC-PPU tissue culture team (coordinated by Edwin Allen), the MRC-PPU Reagents and Services antibody and protein purification teams (coordinated by Dr James Hastie), and the MRC-PPU Genotyping team (coordinated by Gail Gilmour).

## Methods

### Cell Culture and Cas9-expressing cell generation

HEK293T, NIH-3T3, A549 and A549 CRISPR knock-out lines for LRRK2 and PPM1H (Berndsen et al., 2019) were cultured in high glucose DMEM supplemented with glutamine, sodium pyruvate and penicillin-streptomycin. All cells were regularly tested for Mycoplasma PCR products using a Lonza Mycoplasma kit. Before the screen, cells were cultured in the presence of plasmocin as prophylaxis against Mycoplasma infection.

Generation of Cas9 expressing 3T3 cells is described in full detail at protocol.io - <u>dx.doi.org/10.17504/protocols.io.eq2ly7wpmlx9/v1</u>. Briefly, NIH-3T3-Flpin cells were from Thermo Fisher. Early passage cells were transduced with lentivirus carrying HA-Cas9 (Addgene). Cells were selected with blasticidin and single cell sorted onto a 96 well plate. After 2 weeks of culture, twenty individual colonies were picked, expanded, and five were analyzed for Cas9 expression and phosphoRab10, LRRK2, and good growth. The two best clones were tested along with a known positive control lentiviral sgRNA (Pusapati et al., 2018), selected with Puromycin and immunoblotted on day 5 to estimate knock-out efficiency.

Validation of genes using pooled knock-outs: Two gRNA sequences of each gene to be validated were cloned in pLenti-guide puro vector as described (Joung et al. 2017). The plasmids were Sanger sequenced and small scale lentivirus prepared. 3T3-Cas9 cells were infected with lentiviruses, selected for 3 days, and immediately used for immunofluorescence microscopy or immunoblotting.

### Isolation of Rab12 Knockout MEFs

Wild type, heterozygous and homozygous Rab12 knock-out mouse embryonic fibroblasts (MEFs) were isolated from littermate matched mouse embryos at day E12.5 resulting from crosses between heterozygous Rab12 KO/WT mice using a protocol described in <u>dx.doi.org/10.17504/protocols.io.eq2ly713qlx9/v1</u>. Genotypes were verified via allelic sequencing and immunoblotting analysis. Cells were cultured in DMEM containing 10% (v/v) FBS, 2 mM L-glutamine, Penicillin-Streptomycin 100U/mL, 1 mM Sodium Pyruvate, and 1X Non-Essential Amino Acid solution (Life Technologies, Gibco™).

### Expanding the sgRNA genome-wide library

The BRIE library from Addgene was expanded according to protocol.io (dx.doi.org/10.17504/protocols.io.8epv5jr9jl1b/v1). Briefly, the DNA library was electroporated into Lucigen Endura Duos bacteria and the cells plated onto large format Luria broth agar plates to obtain single colonies across the plate. These plates were grown for 14h at 37°C and plasmid extracted using a Machery-Nagel mega-prep kit. Expanded library was PCR amplified using Illumina barcoded PCR primers as described on Addgene (https://media.addgene.org/cms/filer_public/61/16/611619f4-0926-4a07-b5c7-e286a8ecf7f5/broadgpp-sequencing-protocol.pdf) and are part of the supplementary file.

PCR products were sequenced with Miseq to confirm uniform distribution of the gRNA sequences across the population. Aliquots of the plasmid library were frozen at -80°C for future use.

### A flow cytometry based genome wide screen

The detailed protocols can be found at dx.doi.org/10.17504/protocols.io.8epv5jr9jl1b/v1, dx.doi.org/10.17504/protocols.io.eq2ly7wpmlx9/v1.

Briefly, the screen was performed maintaining a 300x fold representation of guides in the transduced cells. For ∼79,500 gRNAs, 3T3-Cas9 cells were plated in 20, 15cm dishes at 5 x 10^6^ cells per dish. Lentiviral gRNAs were infected at an MOI of 0.2 (For ∼100 x 10^6^ cells, ∼ 20 x 10^6^ virus particles). After 48h, cells were passed into 60, 15cm dishes with 1µg/ml Puromycin for selection. After 72h, cells in the control plate that did not receive the virus were dead. Puromycin resistant 3T3-Cas9-BRIE cells were pooled and frozen in cryovial aliquots. Four days before the flow cytometry assay, 40 x 10^6^ cells were thawed and plated into 10, 15cm dishes and allowed to attach and grow for 3 days. On the 4th day, cells were trypsinized, resuspended to a cell density of 5 x 10^6^ cells/ml, passed through a 40µm cell strainer and fixed with 3% PFA for 30min, permeabilized with 0.2% Saponin for 30min and stained overnight at 4°C with rabbit anti-phosphoRab10 antibody at 1µg/ml. Cells were then washed and stained with goat anti-rabbit 647 antibody diluted 2µg/ml for 1h at RT. Cells were washed, resuspended to 2 x 10^6^ cells/ml and injected into a Sony SH800 sorter with FSC of 1, FL4 PMT with a gain of 40% and sample pressure maintained at level 6. MLi-2 treated and secondary antibody alone samples were used as negative controls to identify cell population gates. Cells treated with 4µM nigericin for 3h were positive controls for detection of high level of phosphoRab10.

Cells were sorted based on the histogram of Alexa-647 fluorescent signal. The lowest 5% and highest 7.5% signal containing gates were sorted into two 5ml collection tubes until each had at least 2 x 10^6^ cells. To control for total distribution of gRNAs across the population, 10 x 10^6^ unsorted cells were reserved as input sample. This exercise was performed on two independent sorts from two independent stainings. Sorted cells were pelleted and stored at -80°C for genomic DNA isolation.

### Molecular biology

For genomic DNA extraction, frozen cells were thawed, uncrosslinked and genomic DNA (gDNA) extracted according to dx.doi.org/10.17504/protocols.io.eq2lynm9qvx9/v1. All primers used for PCR amplification for next generation sequencing (NGS) were ordered as Polypak cartridges purified from the Protein and Nucleic Acid facility, Stanford University. Those used for cloning were ordered unpurified. Primer sequences can be found in the supplemental files.

Variable sequences were incorporated in forward primer sequences to improve diversity in the NGS run and 8 such primers were pooled in equimolar ratio [Addgene-P5-F (0-8)]. Reverse primers were incorporated with TrueSeq indices. PCR was performed as described in protocol.io https://dx.doi.org/10.17504/protocols.io.8epv5jr9jl1b/v1. Briefly, input plasmid library and each of the genomic DNA libraries were amplified using Titanium-Taq polymerase. PCR products were cleaned up and size selected using Ampure magnetic beads and concentrated by eluting in small volume, quantified with Qubit high sensitivity dsDNA assay and finally amplicon size confirmed on an Agilent Bioanalyzer. Each PCR amplicon library (two replicates each of unsorted, low phosphoRab10 and high phosphoRab10) was mixed at equimolar ratio and sequenced at Novogene Co., California using their 150 x 2 HiSeq platform.

### Analyses and visualization of next generation sequencing data

Raw sequencing reads were mapped to sgRNA sequence guides in the BRIE library using the count_spacer.py script (Joung et al., 2017) which provided the count of each sgRNA in each sample.

For quality control, evenness of the sgRNA representation was visually assessed by plotting the cumulative distribution of sgRNA representation and quantified using the Gini Index. All samples had a Gini Index lower than 0.42. Consistency between replicates was measured using the Spearman correlation of the sgRNA counts. These quality metrics were computed using Python in a Jupyter notebook available on GitHub (https://github.com/PfefferLab/LRRK2_crispr_screen_paper).

sgRNA effect size estimation: The screen data were analyzed using the MAGeCK MLE algorithm (Wei Li et al 2014). For each gene, MAGeCK MLE collapses the effects of individual sgRNAs into a single gene-level effect size (β-score) and p-value, which quantify the gene contribution to Rab10 phosphorylation in either the positive direction (β-score <0, gene knockout decreases phosphoRab10) or negative direction (β-score >0, gene knockout increases phosphoRab10). p-values were corrected for multiple hypothesis testing using the false discovery rate (FDR) method. Genes with an FDR<0.1 were labeled as either positive regulators (β-score <0) or negative regulators (β-score >0). For this analysis, samples corresponding to the high phosphoRab10, low phosphoRab10, and unsorted population were included in the design matrix with effect coefficients of +1, -1 and 0. Thus, the reported beta score captures the tendency of a gene knockout to push the cells in the high phosphoRab10 (β- score > 0) or low phosphoRab10 population (β-score < 0). For effect size normalization, the 1000 non-targeting sgRNAs of the Brie library were used, and p-values were determined using the permutation method with 100 rounds of permutation.

To assay consistency in the effect direction across individual sgRNAs targeting the same positive or negative regulator genes determined by the MLE method, we calculated guide-level log2 fold change in the high GFP population versus low GFP population using the MAGeCK RRA method. For this analysis, sgRNAs with fewer than 100 counts in both the high and low GFP samples were discarded.

As with the MLE method, effect sizes were normalized using the log2 fold change distribution of the non-targeting sgRNAs.

The MAGeCK output files were loaded as data frames in R and processed with dplyr and ggplot to generate volcano plots, rank plots, and sgRNA-level log2 fold change plots. Code used to run MAGeCK and generate each figure is available on GitHub (https://github.com/PfefferLab/LRRK2_crispr_screen_paper).

All primers, gRNAs, and screen results are included as Supplemental Tables 1-3.

### Lentiviral preparation and transduction

Large scale lentiviral preparation for generating pooled lentiviral gRNA libraries was performed according to a modified protocol from Joung et al (2017) and is published on protocol.io (dx.doi.org/10.17504/protocols.io.8epv5jr9jl1b/v1). Briefly, low passage HEK293T cells were transfected with BRIE library along with the packaging plasmids and viral supernatant was collected 48 h (Day2) and 72 h (Day3) post-transfection. These two separate days of supernatants were pooled, filtered through 0.45µm and frozen at -80°C. An aliquot of the frozen virus was used for titration such that <30% of the cells were transduced and showed Puromycin resistance. An estimate of the number of virus particles / µl was made. For small scale preparations of lentiviruses to express individual gRNAs or GFP-tagged Rab GTPases, a standard lentiviral protocol was used as is published in protocol.io (dx.doi.org/10.17504/protocols.io.bp2l61z2zvqe/v1).

### Effect of PPM1H on Rab12 dependent LRRK2 activation

A549 cells were transduced with the relevant virus (GFP, GFP-Rab12, wtPPM1H-mApple, PPM1H H153D-mApple, PPM1H-D288A mApple) and 5µg/ml polybrene. After 72h, cells were either selected for protein expression with Puromycin or sorted for the relevant fluorescent protein expression. Sorted cells were tested for protein expression by immunoblot.

### 293T overexpression assays

Rab specificity of LRRK2 activation upon overexpression:

HEK293 cells were seeded into six-well plates and transiently transfected at 60-70% confluency using polyethylenimine (PEI) transfection reagent. 1 ug of Flag-LRRK2 WT, R1441C, K17/18A R1441G and 0.5 ug of GFP, GFP-Rab8, GFP-Rab10, GFP-Rab12, or GFP-Rab29 and 7.5 ug of PEI were diluted in 200 uL Opti-MEM™ Reduced serum medium (Gibco™) per well. 36 hours after transfection, cells were treated with 200 nM MLi-2 for 2 hours as indicated and lysed in ice-cold lysis buffer. Samples were prepared for immunoblotting analysis as below.

### LRRK2 activation with Site #3 mutants

HEK293 cells were seeded into six-well plates and transiently transfected at 60–70% confluence using polyethylenimine (PEI) transfection reagent with Flag-LRRK2 wildtype or variant plasmids. 2 µg of plasmid and 6 µg of PEI were diluted in 0.5 ml of Opti-MEM™ Reduced serum medium (Gibco™) per single well. For co-overexpression experiments, 1.6 µg of Flag-LRRK2 wildtype or variant plasmids, 0.4 µg of HA-Rab12, HA-Rab29 or HA-empty, and 6 µg of PEI were diluted in 0.5 ml of Opti-MEM™ Reduced serum medium (Gibco™) per single well. Cells were lysed 24 h post-transfection in an ice-cold lysis buffer containing 50 mM Tris–HCl pH 7.4, 1 mM EGTA, 10 mM 2-glycerophosphate, 50 mM sodium fluoride, 5 mM sodium pyrophosphate, 270 mM sucrose, supplemented with 1 µg/ml microcystin-LR, 1 mM sodium orthovanadate, complete EDTA-free protease inhibitor cocktail (Roche), and 1% (v/v) Triton X-100. Lysates were clarified by centrifugation at 15 000 g at 4°C for 15 min and supernatants were quantified by Bradford assay. Detailed methods for cell transfection and cell lysis can be found in <u>dx.doi.org/10.17504/protocols.io.bw4bpgsn and dx.doi.org/10.17504/protocols.io.b5jhq4j6</u>.

### Mice

The Rab12 knock-out mouse strain used for this research project, C57BL/6N- Rab12em1(IMPC)J/Mmucd (RRID:MMRRC_049312-UCD) was obtained from the Mutant Mouse Resource and Research Center (MMRRC) at University of California at Davis, and was donated to the MMRRC by The KOMP Repository, University of California, Davis (originating from Stephen Murray, The Jackson Laboratory). Mice selected for this study were maintained under specific pathogen-free conditions at the University of Dundee (U.K.). All animal studies were ethically reviewed and carried out in accordance with the Animals (Scientific Procedures) Act 1986 and regulations set by the University of Dundee and the U.K. Home Office. Animal studies and breeding were approved by the University of Dundee ethical committee and performed under a U.K. Home Office project license. Mice were housed at an ambient temperature (20–24°C) and humidity (45–55%) and were maintained on a 12 h light/12 h dark cycle, with free access to food and water. For the experiments described in Figure 2 and FIgure 2- Figure Supplement 1, 3-month-old littermate or age-matched mice of the indicated genotypes were injected subcutaneously with vehicle [40% (w/v) (2-hydroxypropyl)-β-cyclodextrin (Sigma–Aldrich #332607)] or MLi-2 dissolved in the vehicle at a 30 mg/kg final dose. Mice were killed by cervical dislocation 2 h following treatment and the collected tissues were rapidly snap frozen in liquid nitrogen.

### Quantitative immunoblotting analysis

Cells - Quantitative immunoblotting analysis to measure levels of proteins were performed according to the protocol.io <u>dx.doi.org/10.17504/protocols.io.bsgrnbv6</u>. Briefly, cell pellets were collected and lysed in lysis buffer (50 mM Tris–HCl pH 7.4, 1 mM EGTA, 10 mM 2-glycerophosphate, 50 mM sodium fluoride, 5 mM sodium pyrophosphate, 270 mM sucrose, supplemented with 1 μg/ml microcystin-LR, 1 mM sodium orthovanadate, complete EDTA-free protease inhibitor cocktail (Roche), and 1% (v/v) Triton X-100). Lysates were clarified by centrifugation at 10 000 g at 4°C for 10 min. Protein concentration was measured by Bradford and samples equalized and SDS sample buffer added.

Samples were run on 4-20% precast gels (Bio Rad) and transferred onto nitrocellulose membranes. Membranes were blocked in 5% milk with TBST for 1 hour and incubated with specific primary antibodies overnight at 4°C.

Tissues - Quantitative immunoblotting analysis to measure levels of Rab10, phosphoRab10, LRRK2, pS935 LRRK2, and Rab12 was performed as described in <u>dx.doi.org/10.17504/protocols.io.bsgrnbv6</u>. Briefly, snap frozen tissues were thawed on ice in a 10-fold volume excess of ice-cold lysis buffer containing 50 mM Tris–HCl pH 7.4, 1 mM EGTA, 10 mM 2-glycerophosphate, 50 mM sodium fluoride, 5 mM sodium pyrophosphate, 270 mM sucrose, supplemented with 1 μg/ml microcystin-LR, 1 mM sodium orthovanadate, complete EDTA-free protease inhibitor cocktail (Roche), and 1% (v/v) Triton X- 100 and homogenized using a Precellys Evolution system, employing three cycles of 20 s homogenization (6800rpm) with 30 s intervals. Lysates were centrifuged at 15,000g for 30 min at 4°C and supernatants were collected for subsequent Bradford assay and immunoblot analysis.

For blots, primary antibodies used were: Mouse anti-total LRRK2 (Neuromab N241A/34), Rabbit anti- LRRK2 pS935 (UDD2 10(12), MRC PPU Reagents and Services, or ab133450, Abcam), Rabbit anti- LRRK2 pS1292 (ab203181, Abcam), Rabbit anti-pT73 Rab10 (ab230261, Abcam), Mouse anti-total Rab10 (0680–100/Rab10-605B11, Nanotools or ab104859, Abcam), Rabbit anti-pS106 Rab12 (ab256487, Abcam), Rabbit anti-total Rab12 (18843-1-AP, Proteintech), Sheep anti-total Rab12 (SA227, MRC Reagents and Services), Rabbit anti-pS72 Rab7A (ab302494, Abcam), Mouse anti-total Rab7A (R8779, Sigma), Rabbit anti-pT71 Rab29 (ab241062, Abcam), Mouse anti-alpha tubulin (Cell Signaling Technologies, 3873S), Rat anti-HA tag (cat #11867423001, Roche), Sheep anti-PPM1H (DA018, MRC Reagents and Services). Primary antibody probes were detected using IRdye labeled 1:10,000 diluted secondary antibodies (goat anti-mouse 680, goat anti-rabbit 800, goat anti-chicken 680, donkey anti-goat 800). Membranes were scanned on the Licor Odyssey Dlx scanner. Images were saved as .tif files and analyzed using the gel scanning plugin in ImageJ. All primary immunoblotting data associated with each figure has been deposited in a repository and can be found at doi.org/10.5281/zenodo.7633918 and XXXXXXX.

### Immunofluorescence, microscopy and Image analysis

For individual gene knock out validation by microscopy, 3T3-Cas9 cells were transduced with sgRNA lentiviruses for 48 hr, then selected for 3 days with 1µg/ml Puromycin. On Day 6, cells were plated at 30% confluency (75,000 cells) on coverslips in a 24 well plate. After 24h, cells were washed and fixed with 3% paraformaldehyde for 30 min at room temperature, permeabilized with 0.1% Saponin for 30 min, blocked with 2% BSA and stained with rabbit anti-phosphoRab10 and mouse anti-p115 polyclonal antibody for 2 h at room temperature. A549 cells stably expressing GFP-Rab12 and PPM1H-mApple were co-plated with parental A549 cells on coverslips for 24 hrs. Cells were then fixed, stained and imaged for phosphoRab10 as described below.

Cells were washed and stained with DAPI (0.1µg/ml), donkey anti-mouse 488 and donkey anti-rabbit 568 (1:2000) for 1 h at RT. After washing the secondary antibody, coverslips from all wells were mounted on slides using Mowiol. Staining of cells for immunofluorescence is described in the protocol <u>dx.doi.org/10.17504/protocols.io.ewov1nmzkgr2/v1</u>. After the coverslips dried, unbiased multi-position images were obtained using a spinning disk confocal microscope (Yokogawa) with an electron multiplying charge coupled device (EMCCD) camera (Andor, UK) and a 100 × 1.4 NA oil immersion objective. Image acquisition was performed using the multidimensional acquisition using Metamorph. All images were analyzed using an automated pipeline built using Cell Profiler. Whole cell intensities of phosphoRab10 were extracted as median and mean intensities of phosphoRab10 across the cell.

Given the non-uniform nature of the phosphoRab10 dispersal inside cells, median intensity across cell was used for plotting graphs. Images histograms were adjusted on Fiji (https://fiji.sc/) and are presented as maximum intensity projections.

Figures were made in Adobe illustrator. Graphs and statistical analyses were performed in Graphpad Prism.

### LRRK2 Armadillo domain and Rab12 purification

His-Rab12 Q101L, His-LRRK2 Armadillo K439E and E240R were purified after expression in E. coli BL21 (DE3 pLys). Detailed protocols can be found in Gomez et al., 2020 (https://dx.doi.org/10.17504/protocols.io.bffrjjm6) and Vides and Pfeffer, 2021 (https://dx.doi.org/10.17504/protocols.io.bvvmn646). Bacterial cells were grown at 37°C in Luria Broth and induced at A600 nm = 0.6–0.7 by the addition of 0.3 mM isopropyl-1-thio-β-d-galactopyranoside (Gold Biotechnology) and harvested after growth for 18 hr at 18°C. The cell pellets were resuspended in ice-cold lysis buffer (50 mM HEPES, pH 8.0, 10% [vol/vol] glycerol, 500 mM NaCl, 10 mM imidazole, 5 mM MgCl_2_, 0.2 mM tris(2-carboxyethyl) phosphine [TCEP], 20 μM GTP, and EDTA-free protease inhibitor cocktail [Roche]). The resuspended bacteria were lysed by one passage through an Emulsiflex-C5 apparatus (Avestin) at 10,000 lbs/in2 and centrifuged at 40,000 rpm for 45 min at 4°C in a Beckman Ti45 rotor. Cleared lysate was filtered through a 0.2 µm filter (Nalgene) and passed over a HiTrap TALON crude 1 mL column (Cytiva). The column was washed with lysis buffer until absorbance values reached pre-lysate values. Protein was eluted with a gradient from 20 to 500 mM imidazole containing lysis buffer. Peak fractions analyzed by 4-20% SDS-PAGE to locate protein. The eluate was buffer exchanged and further purified by gel filtration on Superdex-75 (GE Healthcare) with a buffer containing 50 mM HEPES, pH 8, 5% (vol/vol) glycerol, 150 mM NaCl, 5 mM MgCl_2_, 0.2 mM tris(2- carboxyethyl) phosphine (TCEP), and 20 μM GTP.

### Microscale Thermophoresis

A detailed method can be found at https://dx.doi.org/10.17504/protocols.io.bvvmn646.

Protein–protein interactions were monitored by microscale thermophoresis using a Monolith NT.115 instrument (NanoTemper Technologies). His LRRK2 Armadillo (1–552) K439E and E240R were labeled using RED-NHS 2nd Generation (Amine Reactive) Protein Labeling Kit (NanoTemper Technologies). For all experiments, unlabeled Rab12 was titrated against a fixed concentration of the fluorescently labeled LRRK2 Armadillo (100 nM); 16 serially diluted titrations of the unlabeled protein partner were prepared to generate one complete binding isotherm. Binding was carried out in reaction buffer (50 mM HEPES pH 8, 150 mM NaCl, 5 mM MgCl_2_, 0.2 mM tris(2-carboxyethyl) phosphine [TCEP], 20 μM GTP, 5% (vol/vol) glycerol, 5 μM BSA, 0.01% Triton-X) in 0.5 mL Protein LoBind tubes (Eppendorf) and allowed to incubate in the dark for 30 min before loading into NT.115 premium treated capillaries (NanoTemper Technologies). A red LED at 20% excitation power (red filter, excitation 605– 645 nm, emission 680–685 nm) and IR-laser power at 60% was used for 30 s followed by 5s of cooling. Data analysis was performed with NTAffinityAnalysis software (NanoTemper Technologies) in which the binding isotherms were derived from the raw fluorescence data and then fitted with both NanoTemper software and GraphPad Prism to determine the Kd using a nonlinear regression method. The binding affinities determined by the two methods were similar. Shown are averaged curves of Rab GTPase-binding partners from two independent experiments, with averaged replicates from each run.

## Figure Legends

**Figure 1 – Figure Supplement 1.**
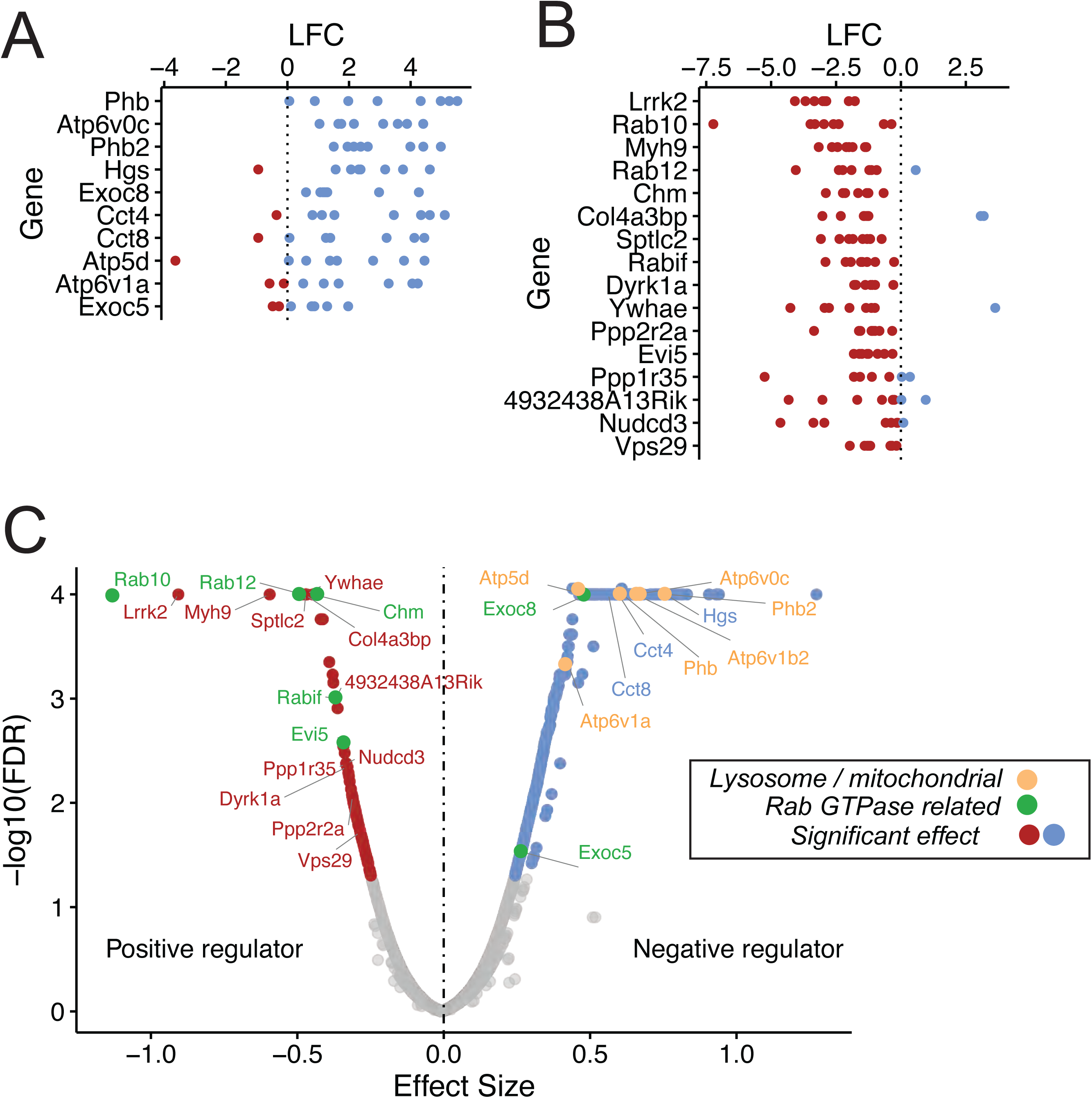
Guide RNA enrichment for CRISPR screen. Log fold change (LFC) in representation of individual guides that target negative regulators **(A)** or positive regulators **(B)**. Each dot represents a single guide; blue and red dots indicate enrichment or de-enrichment in the screen. **(C)** Volcano plot from the MAGeCK MLE analysis; beta score is shown as effect size.

**Figure 1 – Figure Supplement 2.**
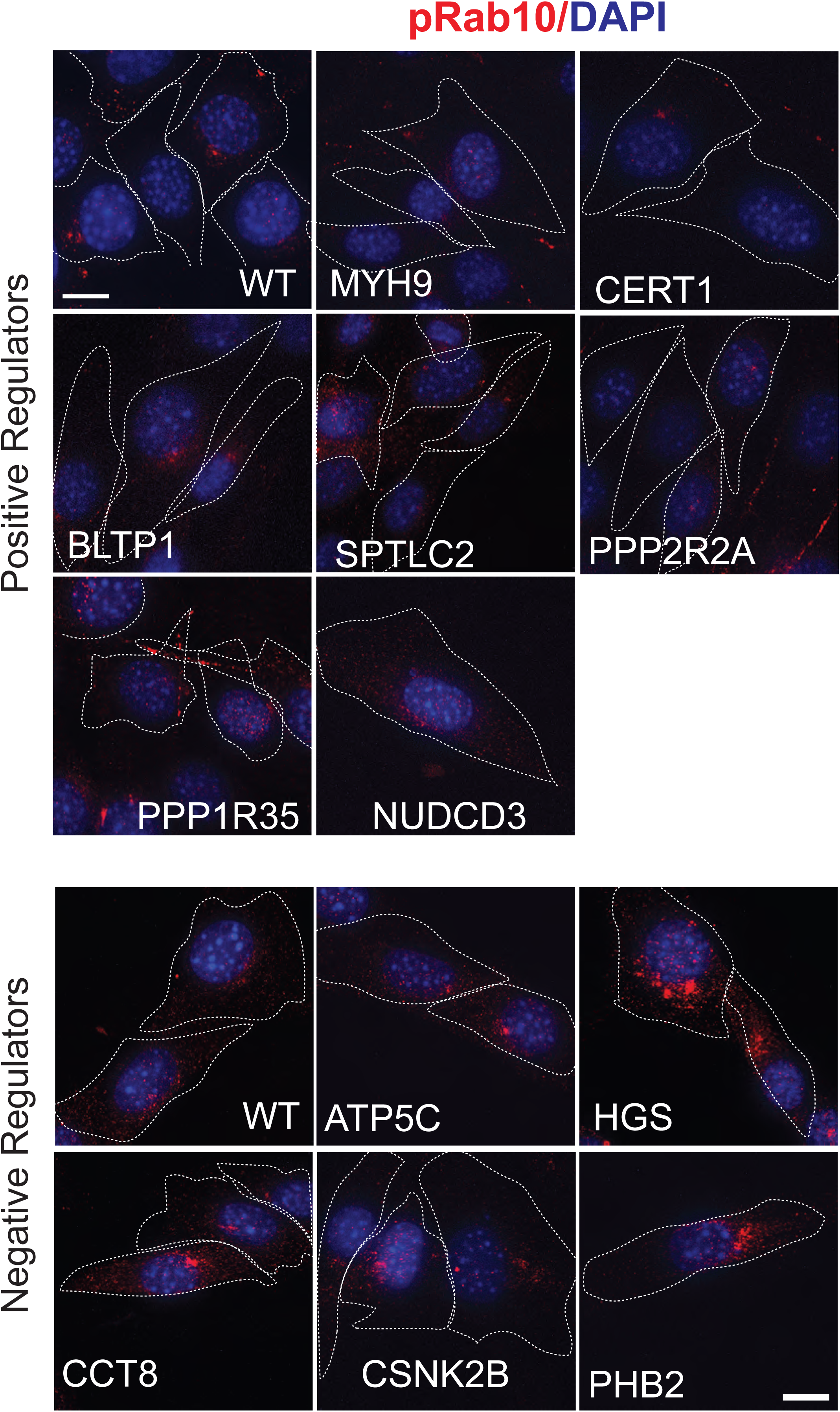
Validation of hits in NIH3T3-Cas9 cells by microscopy. PhosphoRab10was detected by immunofluorescence microscopy in early passage 3T3-Cas9 cells. These cells express lentivirus transduced sgRNAs against individual genes that were top hits. Three days after Puromycin selection cells were stained with rabbit anti-phosphoRab10 antibody. Genes targeted are indicated. Dotted lines indicate the outline of the cells. Scale bar = 10µm.

**Figure 2-Figure Supplement 1.**
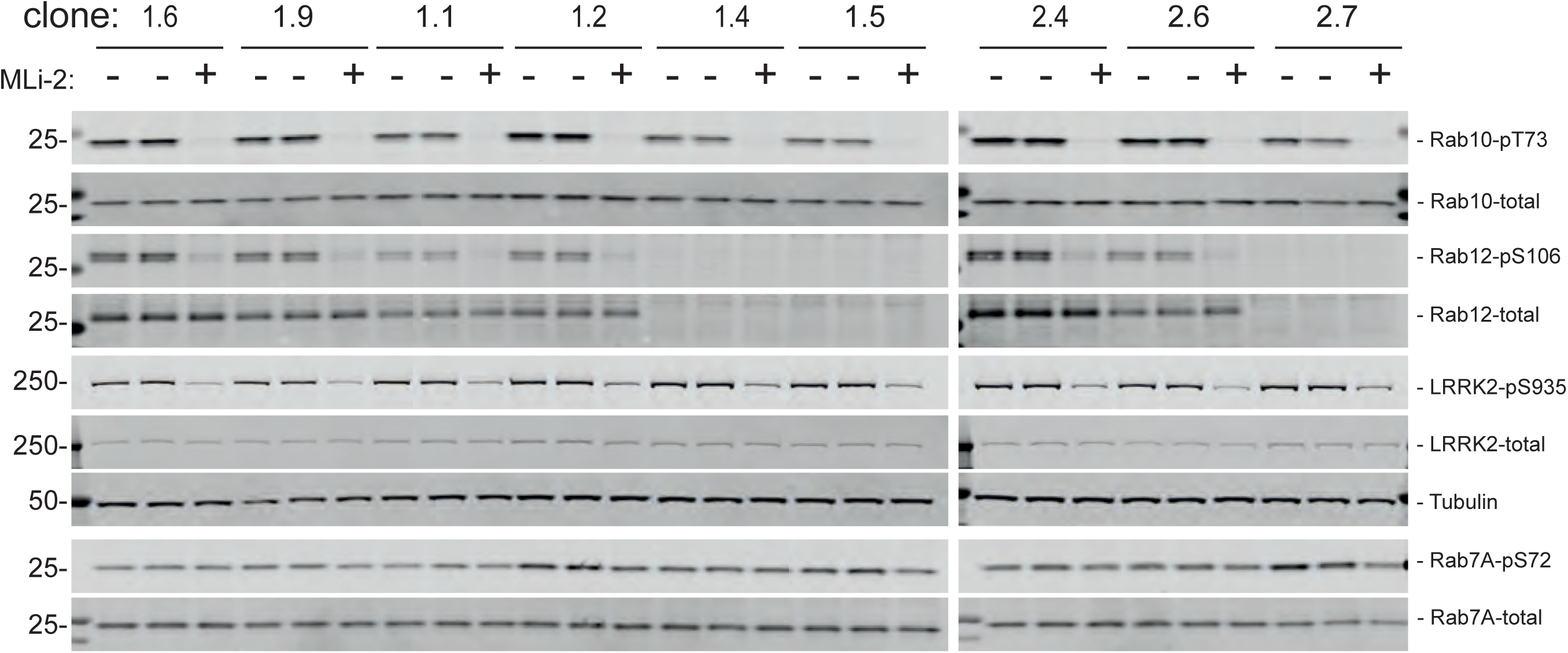
Immunoblots of MEF samples in support of Figure 2.

**Figure 2-Figure Supplement 2.**
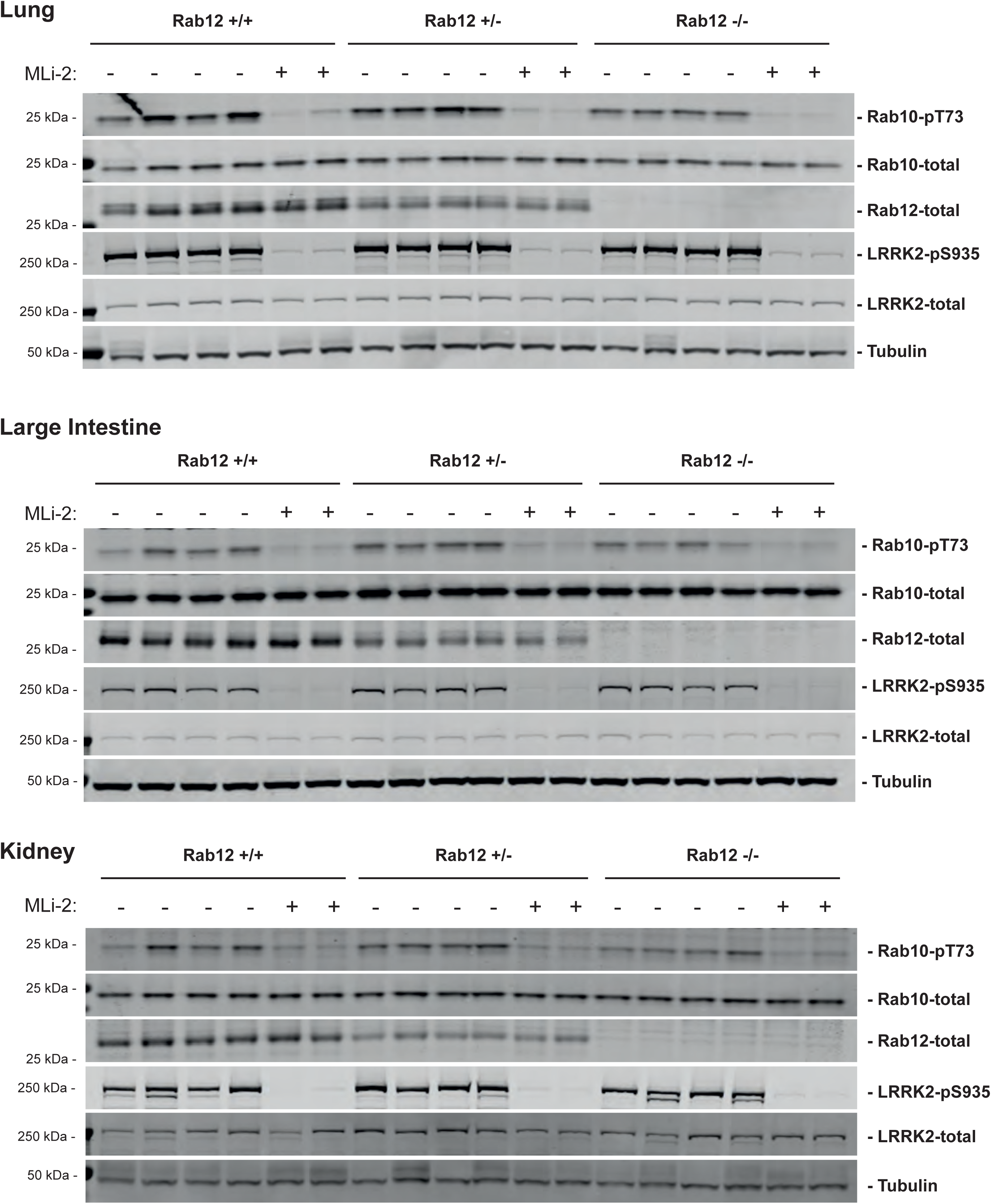
Immunoblots of tissue samples in support of Figure 2.

**Figure 6-Figure Supplement 1.**
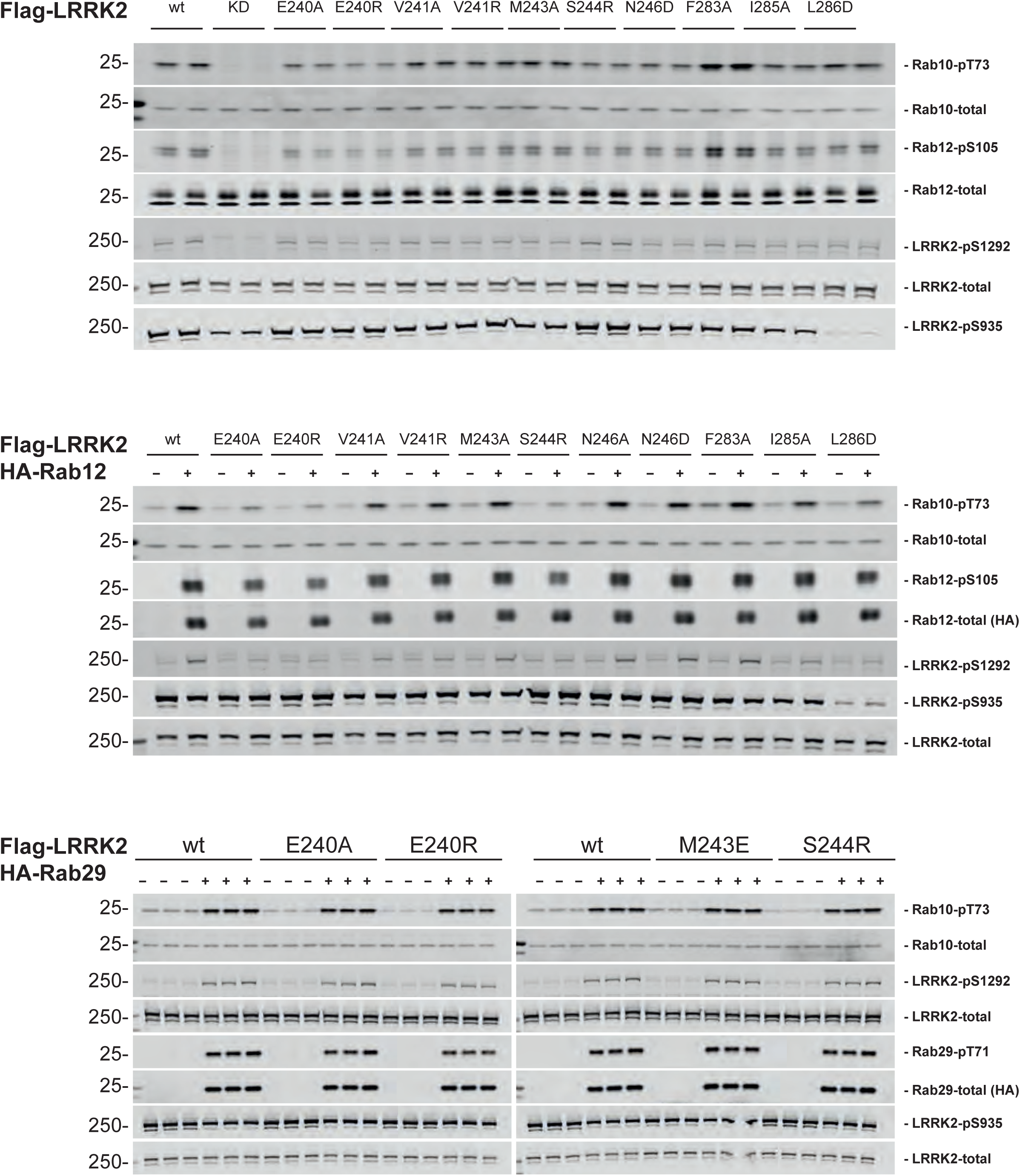
Immunoblots of samples in support of Figure 6.

## Supplemental Tables 1-3.

List of Primers, gRNAs, and all screen results in an Excel File.

